# Molecular evolutionary trends and biosynthesis pathways in the Oribatida revealed by the genome of *Archegozetes longisetosus*

**DOI:** 10.1101/2020.12.10.420141

**Authors:** Adrian Brückner, Austen A. Barnett, Igor A. Antoshechkin, Sheila A. Kitchen

## Abstract

Oribatid mites are a specious order of microarthropods within the subphylum Chelicerata, compromising about 11,000 described species. They are ubiquitously distributed across different microhabitats in all terrestrial ecosystems around the world and were among the first animals colonizing terrestrial habitats as decomposers and scavengers. Despite their species richness and ecological importance genomic resources are lacking for oribatids. Here, we present a 190-Mb genome assembly of the clonal, all-female oribatid mite species *Archegozetes longisetosus* Aoki, a model species used by numerous laboratories for the past 30 years. Comparative genomic and transcriptional analyses revealed patterns of reduced body segmentation and loss of segmental identity gene *abd-A* within Acariformes, and unexpected expression of key eye development genes in these eyeless mites across developmental stages. Consistent with their soil dwelling lifestyle, investigation of the sensory genes revealed a species-specific expansion of gustatory receptors, the largest chemoreceptor family in the genome used in olfaction, and evidence of horizontally transferred enzymes used in cell wall degradation of plant and fungal matter, both components of the *A. longisetosus* diet. Oribatid mites are also noted for their biosynthesis capacities and biochemical diversity. Using biochemical and genomic data, we were able to delineate the backbone biosynthesis of monoterpenes, an important class of compounds found in the major exocrine gland system of Oribatida – the oil glands. Given the mite’s strength as an experimental model, the new high-quality resources provided here will serve as the foundation for molecular research in Oribatida and will enable a broader understanding of chelicerate evolution.

## Introduction

In the past couple of years, the number of sequenced animal genomes has increased dramatically, especially for arthropods about 500 genomes sequences are now available (Childers 2020; Thomas et al. 2020). The majority of these genomes, however, belong to the insects (e.g. flies, beetles, wasp, butterflies and bugs (Thomas et al. 2020)) which compromise the most diverse, yet evolutionarily young and more derived taxa of arthropods (Giribet and Edgecombe 2019; Regier et al. 2010). In strong contrast, genome assemblies, many of which are incomplete or not well annotated, exist for the Chelicerata (Childers 2020) – the other major subphylum of arthropods (Giribet and Edgecombe 2019; Regier et al. 2010). Chelicerates include sea spiders, spiders, mites and scorpions among other organisms, as well as several extinct taxa (Ballesteros and Sharma 2019; Dunlop and Selden 1998). Chelicerates originated as marine animals about 500 million years ago (Dunlop and Selden 1998; Dunlop 2010). Molecular analyses suggest that one particular group, the omnivorous and detritivores acariform mites, may have been among the first arthropods that colonized terrestrial habitats and gave rise to ancient, simple terrestrial food webs (Dunlop and Alberti 2008; Schaefer et al. 2010; Walter and Proctor 1999).

So far, the well-annotated genomic data of chelicerates is limited to animal parasites (including human pathogens and ticks), plant parasites, and predatory mites used in pest control (Cornman et al. 2010; Dong et al. 2017; Dong et al. 2018; Grbić et al. 2011; Gulia-Nuss et al. 2016; Hoy et al. 2016; Rider et al. 2015). Other than some lower-quality genome assemblies (Bast et al. 2016), there are no resources available for free-living soil and litter inhabiting species. Such data are, however, pivotal to understanding the evolution of parasitic lifestyles from a free-living condition and to bridge the gap between early aquatic chelicerates such as horseshoe crabs, and highly derived terrestrial pest species and parasites (Klimov and OConnor 2013; Shingate et al. 2020; Weinstein and Kuris 2016). Because the phylogeny of Chelicerata remains unresolved, additional chelicerate genomes are urgently needed for comparative analyses (Ballesteros and Sharma 2019; Dunlop 2010; Lozano-Fernandez et al. 2019). To help address this deficit, we report here the genome assembly of the soil dwelling oribatid mite *Archegozetes longisetosus* (Aoki, 1965; Figure 1) (Aoki 1965) and a comprehensive analysis in the context of developmental genes, feeding biology, horizontal gene transfer and biochemical pathway evolution of chelicerates.

**Figure 1.**
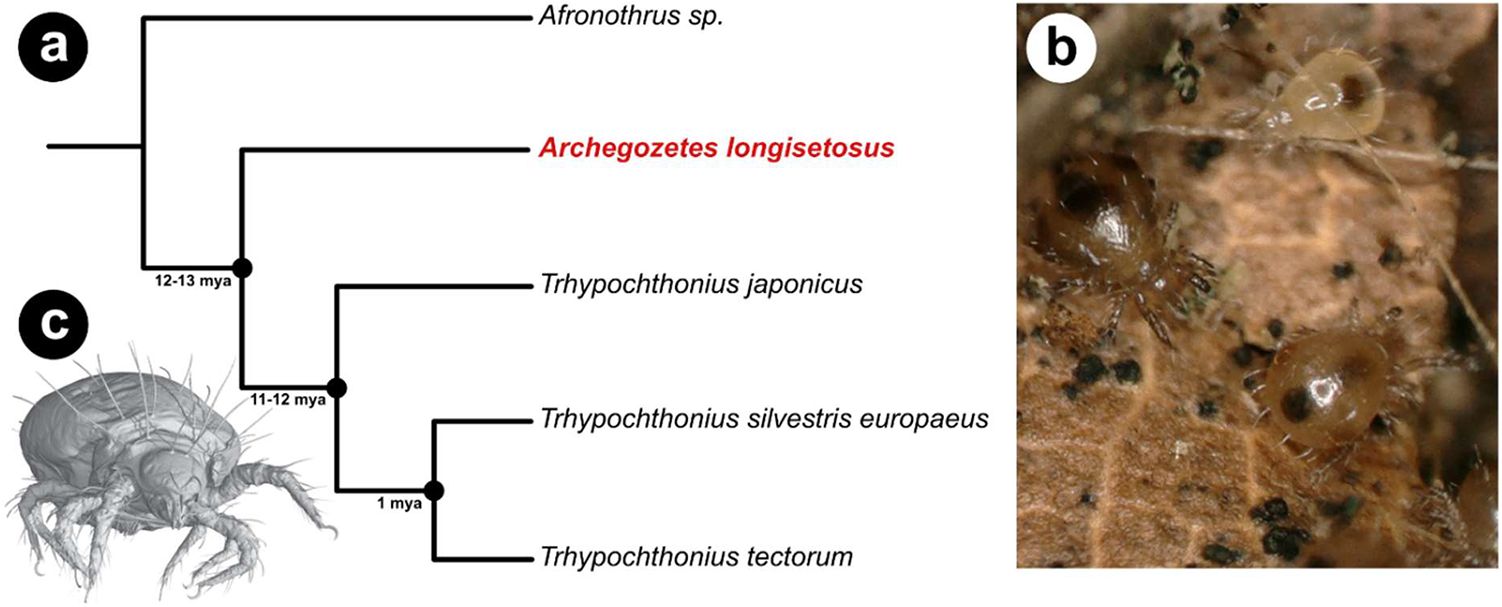
The mite, *Archegozetes longisetosus*, in its phylogenetic and natural environment. a: Species tree of selected oribatid mites of the family Trhypochthoniidae based on phylogenetic analyses and divergence time estimates by (Heethoff et al. 2011b). b: Two adults and one tritonymph of *Archegozetes* on a piece of leaf litter. The algae growing on the leaf serves as a food source for the mites. c: Habitus of an adult mite based on a surface rendering of a µCT-scan reconstruction. Image courtesy of Sebastian Schmelzle.

*Archegozetes longisetosus* (hereafter referred to as *Archegozetes*) is a member of the Oribatida (Acariformes, Sarcoptiformes), an order of chelicerates well-known for their exceptional biosynthesis capacities, biochemical diversity, unusual mode of reproduction, unusually high pulling strength, mechanical resistances and pivotal ecological importance (Brückner et al. 2020; Brückner et al. 2017b; Heethoff and Koerner 2007; Heethoff et al. 2009; Maraun et al. 2007; Maraun and Scheu 2000; Norton and Palmer 1991; Raspotnig 2009; Schmelzle and Blüthgen 2019). *Archegozetes*, like all members of its family Trhypochthoniidae (Figure 1a), reproduce via thelytoky (Heethoff et al. 2013). That means the all-female lineages procreate via automictic parthenogenesis with an inverted meiosis of the holokinetic chromosomes, resulting in clonal offspring (Bergmann et al. 2018; Heethoff et al. 2006; Palmer and Norton 1992; Wrensch et al. 1994). While studying a parthenogenetic species is useful for the development of genetic tools as stable germ-line modifications can be obtained from the clonal progeny without laboratory crosses, one is confronted with the technical and philosophical problems of species delineation, cryptic diversity and uncertain species distribution (Heethoff et al. 2013; Oxley et al. 2014). Reviewing all available data, Norton (Norton 1994; 2007) and Heethoff et al. (Heethoff et al. 2013) concluded that *Archegozetes* is found widely on continents and islands throughout the tropical and partly subtropical regions of the world and that it is a middle-derived oribatid mite closely related to the suborder Astigmata.

One major feature of most oribatid mites is a pair of opisthonotal oil-glands and *Archegozetes* is no exception (Raspotnig 2009; Sakata and Norton 2001). These are a pair of large exocrine glands, each composed of a single-cell layer invagination of the cuticle, which is the simplest possible paradigm of an animal gland (Brückner and Parker 2020; Heethoff 2012). The biological role of these glands was rather speculative for a long time; ideas ranged from a lubricating and osmo- or thermoregulative function (Riha 1951; Smrž 1992; Zachvatkin 1941) to roles in chemical communication (Heethoff et al. 2011a; Raspotnig 2006; Shimano et al. 2002). So far about 150 different gland components have been identified from oribatid mites, including mono- and sesquiterpenes, aldehydes, esters, aromatics, short-chained hydrocarbons, hydrogen cyanide (HCN) and alkaloids (Brückner et al. 2017b; Brückner et al. 2015; Heethoff et al. 2018; Raspotnig 2009; Saporito et al. 2007). While some chemicals appear to be alarm pheromones (Raspotnig 2006; Shimano et al. 2002), most function as defensive allomones (Heethoff et al. 2011a). Interestingly, alkaloids produced by oribatids mites are the ultimate source of most toxins sequestered by poison-frogs (Saporito et al. 2007; Saporito et al. 2009).

Terrestrial chelicerates predominately ingest fluid food. While phloem-feeding plant pests like spider mites and ecotoparasites likes ticks adapted a sucking feeding mode, scorpions, spiders and others use external, pre-oral digestion before ingestion by morphologically diverse mouthparts (Bensoussan et al. 2016; Cohen 1995; Dunlop and Alberti 2008; Gulia-Nuss et al. 2016). Exceptions from this are Opiliones and sarcoptiform mites, i.e. oribatid and astigmatid mites, all of which ingest solid food (Heethoff and Norton 2009; Norton 2007; Shultz 2007). In general, oribatids feed on a wide range of different resources and show a low degree of dietary specialization (Brückner et al. 2018b). The typical food spectrum of Oribatida, includes leaf-litter, algae, fungi, lichens, nematodes, and small dead arthropods such as collembolans (Heidemann et al. 2011; Riha 1951; Schneider and Maraun 2005; Schneider et al. 2004a; Schneider et al. 2004b). In laboratory feeding trials, oribatid mites tend to prefer dark pigmented fungi, but also fatty acid-rich plant-based food (Brückner et al. 2018b; Schneider and Maraun 2005). Additionally, stable-isotope analyses of ^15^N and ^13^C suggested that Oribatida are primary- and secondary decomposers feeding on dead plant material and fungi, respectively (Maraun et al. 2011; Schneider et al. 2004a). The reasons for these preferences are still unknown, but they raise the question of how oribatid mites are able to enzymatically digest the cell walls of plants and fungi (Brückner et al. 2018a; Brückner et al. 2018b; Schneider et al. 2004b; Smrž and Čatská 2010).

Early studies on *Archegozetes* and other mites found evidence for cellulase, chitinase and trehalase activity which was later attributed to symbiotic gut bacteria (Haq 1993; Luxton 1979; Siepel and de Ruiter-Dijkman 1993; Smrž 2000; Smrž and Čatská 2010; Smrž and Norton 2004; Zinkler 1971). While such bacterial symbionts are a possible explanation, genomic data of other soil organisms and plant-feeding arthropods suggest a high frequency of horizontal transfer of bacterial and fungal genes enabling the digestion of cell walls (Grbić et al. 2011; Mayer et al. 2011; McKenna et al. 2019; Wu et al. 2017; Wybouw et al. 2016; Wybouw et al. 2018). For instance, an in-depth analysis of the spider mite *Tetranychus urticae* revealed a massive incorporation of microbial genes into the mite’s genome (Grbić et al. 2011; Wybouw et al. 2018). Horizontal gene transfer appears to be a common mechanism for soil organisms, including mites, to acquire novel metabolic enzymes (Dong et al. 2018; Faddeeva-Vakhrusheva et al. 2016; Grbić et al. 2011; Hoffmann et al. 1998; Mayer et al. 2011; Wu et al. 2017), and hence seems very likely for *Archegozetes* and other oribatid mite species that feed on plant or fungal matter.

*Archegozetes* has been established as a laboratory model organism for three decades, having been used in studies, ranging from ecology, morphology, development and ecotoxicology to physiology and biochemistry (Barnett and Thomas 2012; 2013a; 2013b; 2018; Brückner et al. 2017a; Brückner et al. 2020; Heethoff et al. 2013). As such, *Archegozetes* is among the few experimentally tractable soil organisms and by far the best-studied oribatid mite species (Barnett and Thomas 2012; Heethoff et al. 2013; Thomas 2002). Since it meets the most desirable requirements for model organisms (Thomas 2002), that is a rapid development under laboratory conditions, a dedicated laboratory strain was named *Archegozetes longisetosus* ran in reference to its founder Roy A. Norton (Heethoff et al. 2013, Figure 1b-c). Their large number of offspring enables mass cultures of hundreds of thousands of individuals, and their cuticular transparency during juvenile stages, and weak sclerotization as adults are general assets of an amenable model system (Brückner et al. 2018c; Brückner et al. 2016; Heethoff et al. 2013; Heethoff and Raspotnig 2012). In the past 10 years, *Archegozetes* also received attention as a model system for chemical ecology (Brückner and Heethoff 2018; Brückner et al. 2020; Brückner et al. 2016; Heethoff and Rall 2015; Heethoff and Raspotnig 2012; Raspotnig et al. 2011; Thiel et al. 2018). Some of these studies focusing on the *Archegozetes* gland revealed basic insights into the chemical ecology and biochemical capabilities of arthropods (Brückner et al. 2020; Heethoff and Rall 2015; Thiel et al. 2018). Hence, *Archegozetes* is poised to become a genetically tractable model to study the molecular basis of gland and metabolic biology.

The aim and focus of the current study were three-fold – to provide well-annotated, high-quality genomic and transcriptomic resources for *Archegozetes longisetosus* (Figure 1), to reveal possible horizontal gene transfers that could further explain the feeding biology of oribatids, and to present *Archegozetes* as a research model for biochemical pathway evolution. Through a combination of comparative genomic and detailed computational analyses, we were able to generate a comprehensive genome of *Archegozetes* and provide it as an open resource for genomic, developmental and evolutionary research. We further identified candidate horizontal gene transfer events from bacteria and fungi that are mainly related to carbohydrate metabolism and cellulose digestion, features correlated with the mite feeding biology. We also used the genomic data together with stable-isotope labeling experiments and mass spectrometric investigation to delineate the biosynthesis pathway of monoterpenes in oribatid mites.

## Results and Discussion

### *Archegozetes longisetosus* genome assembly

*Archegozetes longisetosus* (Figure 1) has a diploid chromosome number (2n) of 18 (Heethoff et al. 2006), most likely comprising 9 autosomal pairs, the typical number of nearly all studied oribatid mite species (Norton et al. 1993). There are no distinct sex chromosomes in *Archegozetes*; this appears to be ancestral in the Acariformes and persisted in the Oribatida (Heethoff et al. 2006; Norton et al. 1993; Wrensch et al. 1994). Even though some XX:XO and XX:XY genetic systems have been described in the closely related Astigmata, the sex determination mechanism in oribatids, including *Archegozetes*, remains unknown (Heethoff et al. 2013; Heethoff et al. 2006; Norton et al. 1993; Oliver Jr 1983; Wrensch et al. 1994). To provide genetic resources, we sequenced and assembled the genome using both Illumina short-read and Nanopore MinION long-read sequencing approaches (**Table 1**; see also “**Materials and Methods**”). Analyses of the *k-mer* frequency distribution of short reads (Table 1; Supplementary Figure S1) resulted in an estimated genome size range of 135-180 Mb, smaller than the final assembled size of 190 Mb (**Table 1**; see also “**Materials and Methods**”). This difference was suggestive of high repetitive content in the genome of *Archegozetes* and indeed, repeat content was predicted to be 32 % of the genome (see below) (Alfsnes et al. 2017; Simpson 2014). Compared to genome assemblies of other acariform mites, the assembled genome size of *Archegozetes* is on the large end, but is smaller than that of mesostigmatid mites, ticks and spiders (Bast et al. 2016; Dong et al. 2017; Dong et al. 2018; Grbić et al. 2011; Gulia-Nuss et al. 2016; Hoy et al. 2016; Schwager et al. 2017). In the context of arthropods in general, *Archegozetes*’s genome (**Table 1**) is among the smaller ones and shares this feature with other arthropod model species like the spider mite, *Drosophila*, clonal raider ant and red flour beetle (Consortium 2008; dos Santos et al. 2015; Grbić et al. 2011; Oxley et al. 2014). Even though we surface-washed the mites and only used specimens with empty alimentary tracts for sequencing, we removed 438 contigs with high bacterial or fugal homology making up approximately 8.5 Mb of contamination (s**ee supplementary Table S1**). The final filtered genome assembly was composed of 1182 contigs with an N_50_ contiguity of 994.5 kb (**Table 1**).

**Table 1.** *Archegozetes longisetosus* genome metrics

| Feature | Value |
| --- | --- |
| Estimated genome size | 135-180 Mb |
| Assembly size | 190 Mb |
| Coverage based on assembly (short/long) | 200x (short), ~60x (long) |
| # contigs | 1182 |
| N50 (contigs) | 994.5 kb |
| Median contig length | 50.3 kb |
| GC content | 30.9% |
| # gene models | 23,825 |

### The official gene set and annotation of *Archegozetes*

We generated the official gene set (OGS) for *Archegozetes* by an automated, multi-stage process combining *ab inito* and evidenced-based (RNAseq reads, transcriptomic data and curated protein sequences) gene prediction approaches (see “**Materials and Methods**”) yielding 23,825 gene models. In comparison to other mites and ticks as well as insects, this is well within the range of the numbers discovered in other Chelicerata so far (Figure 2a). Chelicerates with a large OSG, however, usually possess larger genomes (1-7 Gb), which suggests that *Archegozetes* may have a relatively dense distribution of protein-coding genes in its genome. On the other hand, ticks can have giga-base sized genomes, but only a rather small number of gene models, probably due to high repetitive content (Barrero et al. 2017; Gulia-Nuss et al. 2016; Palmer et al. 1994; Van Zee et al. 2007). Lacking more high-quality genomic resources of mites, it is thus not clear whether the OGS of *Archegozetes* is the rule, or rather the exception within the Oribatida.

**Figure 2.**
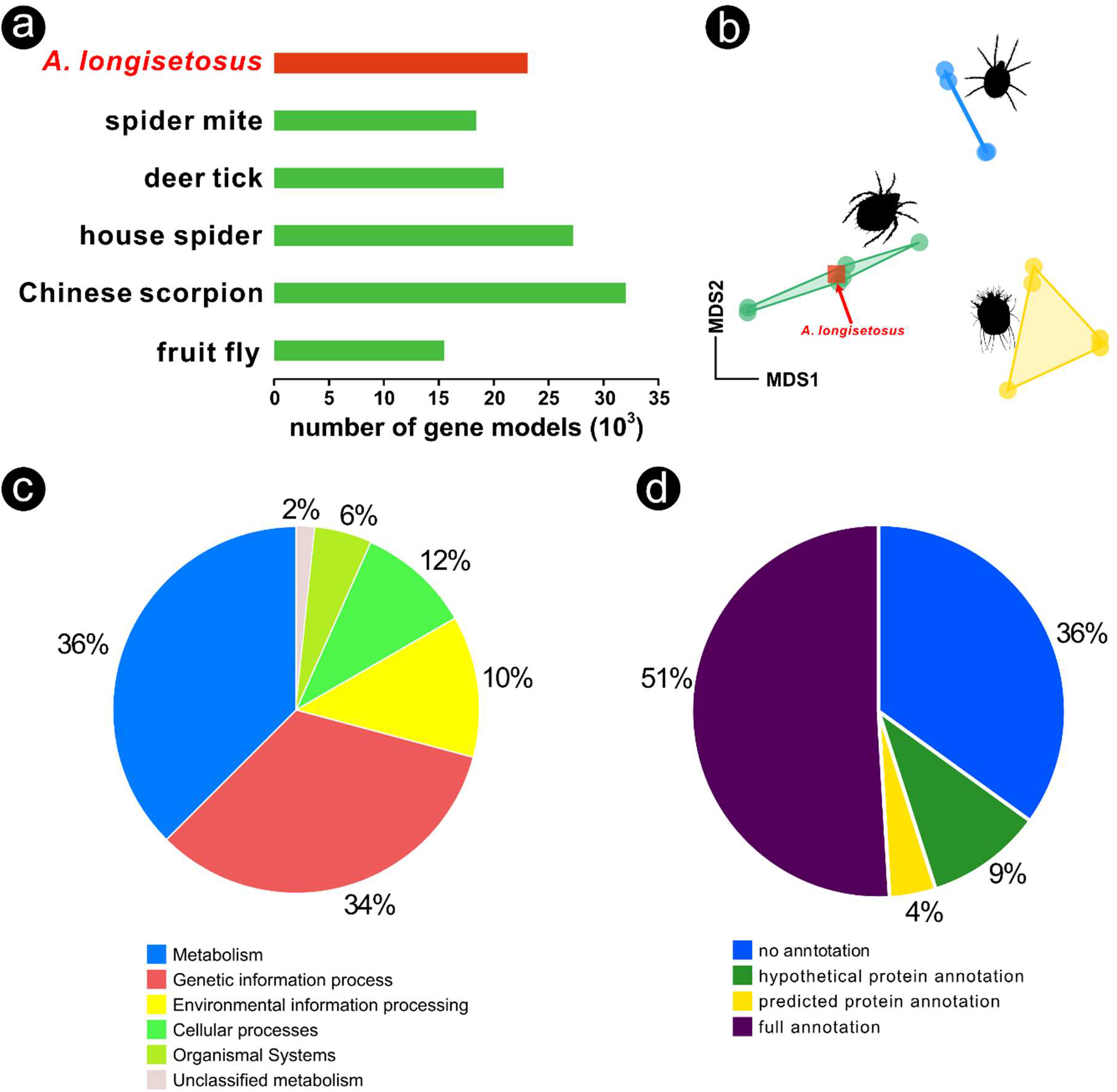
Comparisons and annotations of the official gene set (OGS) of *Archegozetes* longisetosus. a: Number of gene models of the mites compared to other mites, chelicerates and the fruit fly (Cao et al. 2013; dos Santos et al. 2015; Grbić et al. 2011; Gulia-Nuss et al. 2016; Schwager et al. 2017). b: Non-linear multidimensional scaling plot (NMDS) of clustered orthogroups based on the OGS or predicted proteins of several mite species. *Archegozetes longisetosus* is marked a square, nested within Oribatida. Prostigmata are depicted in blue, Astigmata in yellow and Oribatida in green. c: Pie chart showing the percentage composition of genes of the *Archegozetes* annotated to different broad biological categories by GhostKOALA. d: Pie chart describing the overall annotation of the OGS of the mite.

To compare if *Archegozetes*’ OSG is similar to predicted genes of other oribatid mites as well as Prostigmata and Astigmata, we first clustered genes by ortholog inference (OrthoFinder; (Emms and Kelly 2015b)), removed species-specific genes and constructed a presence-absence matrix of orthogroups to ordinate the data using non-metric multidimension scaling (NMDS, Figure 2b). Ordination revealed that the OGS of *Archegozetes* is well nested with other oribatid mites and clearly separated from their closest relative the astigmatid mites as well as prostigmatid mites (Figure 2b). As a first step in annotating the OSG, we ran KOALA (KEGG Orthology And Links Annotation) to functionally characterize the genes (Kanehisa et al. 2016). In total, 10,456 (43.9%) of all genes received annotation and about two thirds of all genes were assigned either as metabolic genes (36%) or genes related to genetic information processing (34%), while the remaining genes fell into different KEGG categories (Figure 2c). To further annotate the genome, we followed the general workflow of funannotate with some modifications (Palmer and Stajich 2017, see “Materials and Methods”).

Overall, we found 15,236 genes (64%) of the OGS with homology to previously published sequences (Figure 2d). For about half of all genes (51%), we were able to assign a full annotation, 4% of all genes only showed homology to bioinformatically predicted proteins of other species, while 9% of all genes only showed homology to hypothetical proteins (Figure 2d). As only a few high-quality, annotated mite genomes are available and the two-spotted spider mite is the sole species with any experimentally confirmed gene models, it is not surprising that we were only able to confidently annotate about 55% of all genes of the OGS (Figure 2d).

### Orthology and comparative genomics of chelicerates

To further access the protein-coding genes of the mite, we compared the OGS to other chelicerates. Both concatenated maximum likelihood and coalescent species-tree phylogenomic approaches based on 1,121 orthologs placed *Archegozetes*, as expected, within the Nothrina (Heethoff et al. 2013; Pachl et al. 2012) with strong support and recovered previously found oribatid clade topologies (Figure 3a). Our analysis placed the Astigmata as a sister group of Oribatida and not nested within oribatids as suggested based on life-history, chemical defensive secretions, morphology and several molecular studies (Alberti and Michalik 2004; Dabert et al. 2010; Domes et al. 2007; Klimov et al. 2018; Koller et al. 2012; Li and Xue 2019; Liana and Witaliński 2005; Maraun et al. 2004; Norton 1994; 1998; Pepato and Klimov 2015; Sakata and Norton 2001). The relationship of Oribatida and Astigmata has been challenging to resolve for the past decades and several studies using different set of genes, ultra-conserved elements or transcriptomic data reconstructed discordant phylogenies, some of which are similar to ours (Dabert et al. 2010; Domes et al. 2007; Klimov et al. 2018; Li and Xue 2019; Lozano-Fernandez et al. 2019; Maraun et al. 2004; Pepato and Klimov 2015; Van Dam et al. 2019). Overall, the Oribatid-Astigmatid relationship remains unresolved and a broader taxon sampling, especially of more basal Astigmata, will be necessary (Domes et al. 2007; Klimov et al. 2018; Lozano-Fernandez et al. 2019; Norton 1994; 1998; Van Dam et al. 2019). We recovered Trombidiformes (Prostigmata and Sphaerolichida) as sister group of the Sarcoptiformes (Oribatida and Astigmata) constituting the Acariformes (Figure 3a). Neither the maximum likelihood phylogeny (Figure 3a), nor the coalescence-based phylogeny (**Supplementary Figure S2**) reconstructed the Acari (i.e. Acariformes and Parasitiformes) as a monophyletic taxon. Even though there is morphological, ultrastructural and molecular evidence for a biphyletic Acari, aswe recovered here, this relationship and larger-scale chelicerate relationships remain unclear (Alberti 1984; 1991; Dabert 2006; Dunlop and Alberti 2008; Jeyaprakash and Hoy 2009; Li and Xue 2019; Lozano-Fernandez et al. 2019; Van Dam et al. 2019).

**Figure 3.**
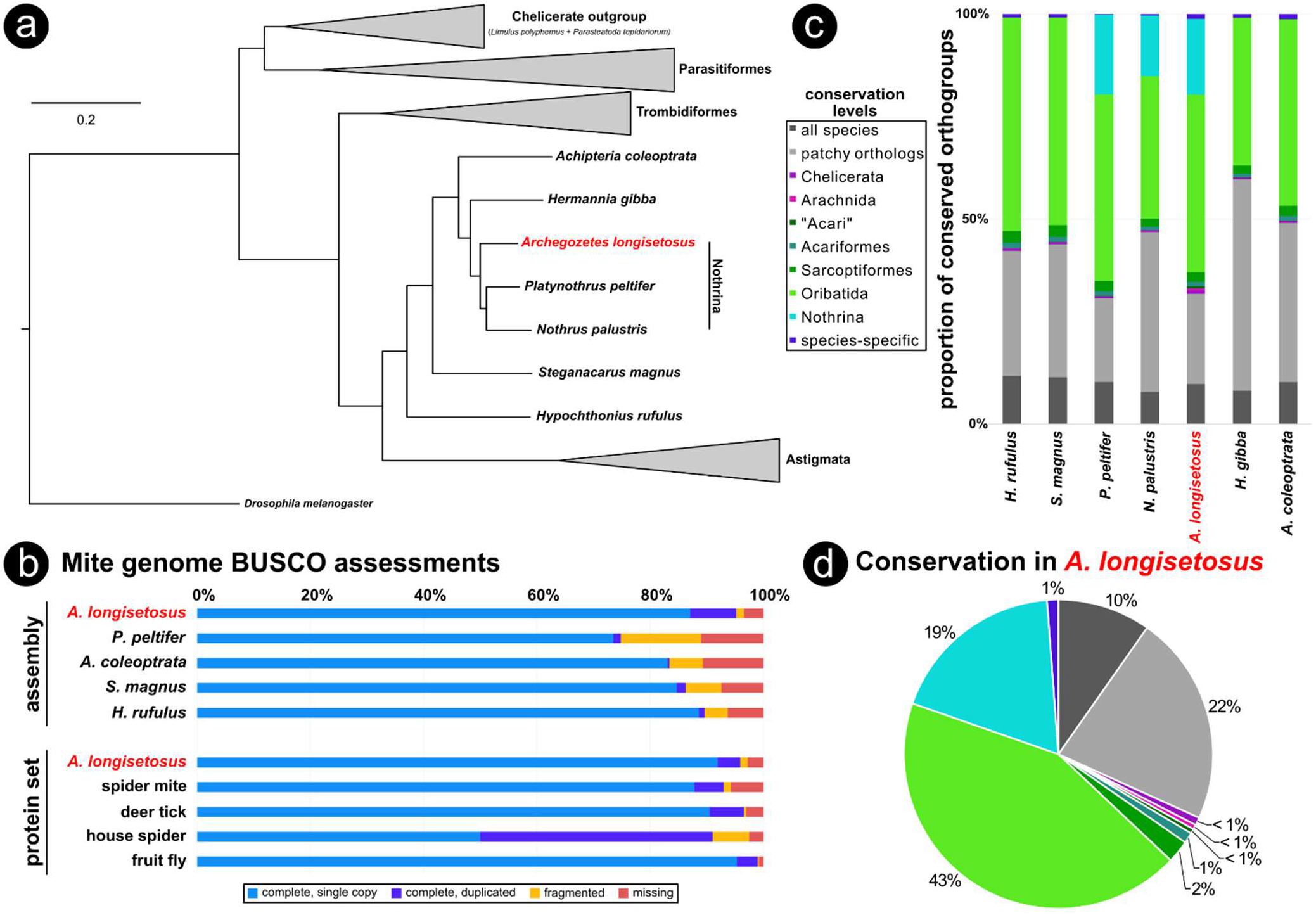
Orthology comparison and phylogenetic placement of *Archegozetes longisetosus* among other chelicerates. a: Maximum likelihood phylogeny based on concatenation of 1,121 orthologs showing the mites phylogenetic environment within the Oribatida (all nodes have 100% support; branch length unit is substitutions per site). For the fully expanded tree see supplementary Figure S2. b: BUSCO-assessment of the *Archegozetes* genome assembly and protein set for both ortholog presence and copy number compared to other oribatid mites and selected model species, respectively. c: Comparisons of protein-coding genes in seven oribatid mite species (for a full comparison of all species see supplementary Figure S3) with *Archegozetes* highlighted in red. The bar charts show the proportion of orthrogroup conservation with each species (see insert legend) based on OrthoFinder clustering. d: Detailed pie chart depicting the conservation levels of orthogroup in *Archegozetes*.

To further assess the quality and homology of both the genome assembly (**Table 1**) and the OGS (Figure 2), we used the1066 arthropod Benchmarking Universal Single-Copy Ortholog (BUSCO) genes data set (Simão et al. 2015). Nearly all BUSCO genes were present in the *Archegozetes* assembly and OGS (96.2% and 97.3%, respectively; Figure 3b). Compared to other genomes sequenced so far, the *Archegozetes* genome has the highest completeness among oribatid mites and the OGS completeness is on par to the well curated genomes of other chelicerate species and *Drosophila melanogaster* (Figure 3b). This result is not surprising because the *Archegozetes* genome was assembled from long-read and short-read data, while all other oribatid mite genomes were solely short reads sequenced on older Illumina platforms (Bast et al. 2016). The fraction of duplicated BUSCO genes in *Archegozetes* (4%) was similar to that of the spider mite and deer tick (Grbić et al. 2011; Gulia-Nuss et al. 2016), but very low compared to the house spider (Figure 3c), whose genome underwent an ancient whole-genome duplication (Schwager et al. 2017).

Overall, the high quality of both the genome assembly and OGS of *Archegozetes* compared to those of other oribatid mites, strongly indicates the importance of this genomic resource. We next categorized all protein models from the OGS by conversation level based on a global clustering orthology analysis (OrthoFinder; Emms and Kelly 2015b) of 23 species (Figure 3c**; supplementary Figure S3**) representing Acariformes, Parasitiformes, several other chelicerates and the fly Drosophila. As for most other species (Siepel et al. 2005; Thomas et al. 2020), about a third of all orthogroups was highly conserved (Figure 3c) across the arthropods, being either in all species (10%; Figure 3d) or is most (22%; Figure 3d). Only 1% of all *Archegozetes* orthogroups did not show homology and were species specific (Figure 3c and d). Only a low proportion (Figure 3c) of orthogroups was conserved across the higher taxonomic levels (all <1% in *Archegozetes*; Figure 3d), which is in line with previous studies that included prostigmatid and mesostigmatid mites (Dong et al. 2017; Dong et al. 2018; Hoy et al. 2016). Interestingly, there was a large proportion of orthogroups conserved across all Oribatida (43% in *Archegozetes*; Figure 3d) and also about 19% of orthogroups in *Archegozetes* were shared only with other Nothrina (Figure 3d). A fairly large percentage of these orthogroups may contain potentially novel genes that await experimental verification and functional analyses (Emms and Kelly 2015b; Nagy et al. 2020; Thomas et al. 2020). Especially the lack of homology within the Sarcoptiformes (2-3%; Figure 3c) may explain the controversial placement of Astigmata as a sistergroup of Oribatida that we recovered (Figure 3a). This grouping is likely caused by a long-branch attraction artifact and the sister relationship was incorrectly inferred (Dabert 2006; Dabert et al. 2010; Domes et al. 2007; Klimov et al. 2018; Pepato and Klimov 2015), because orthogroup clustering could not detect enough homology between oribatids and the Astigmata so far sequenced, which are highly derived. Hence, a broad taxon sampling of basal astigmatid mite genomes seems necessary to resolve Oribatida-Astigmata relationship (Li and Xue 2019; Norton 1994; 1998; Pepato and Klimov 2015; Van Dam et al. 2019).

### Repeat content analysis and transposable elements (TEs)

For clonal species like *Archegozetes*, reproducing in the absence of recombination, it has been hypothesized that a reduced efficacy of selection could results in an accumulation of deleterious mutations and repeats in the genome (Arkhipova and Meselson 2000; Barton 2010; Charlesworth 2012; Muller 1964; Nuzhdin and Petrov 2003; Schön et al. 2009). There is, however, no evidence for such an accumulation in oribatids or other arthropods (Bast et al. 2016). Generally, we found that most of the repetitive content in *Archegozetes* could not be classified (57%; Figure 4a). The high proportion of unknown repeats likely corresponds to novel predicted repetitive content, because of limited repeat annotation of mites in common repeat databases such as RepBase (Bast et al. 2016). Regarding the two major classes of repeat content, DNA transposons made up about 32% of total repeats, while only 5% represented retrotransposons (Figure 4a). About 6% of total repetitive content comprised simple and low complexity repeats (Figure 4a). Overall, the total repetitive content (32%, Figure 4b) seems to be within a normal range for chelicerates and arthropods.

**Figure 4.**
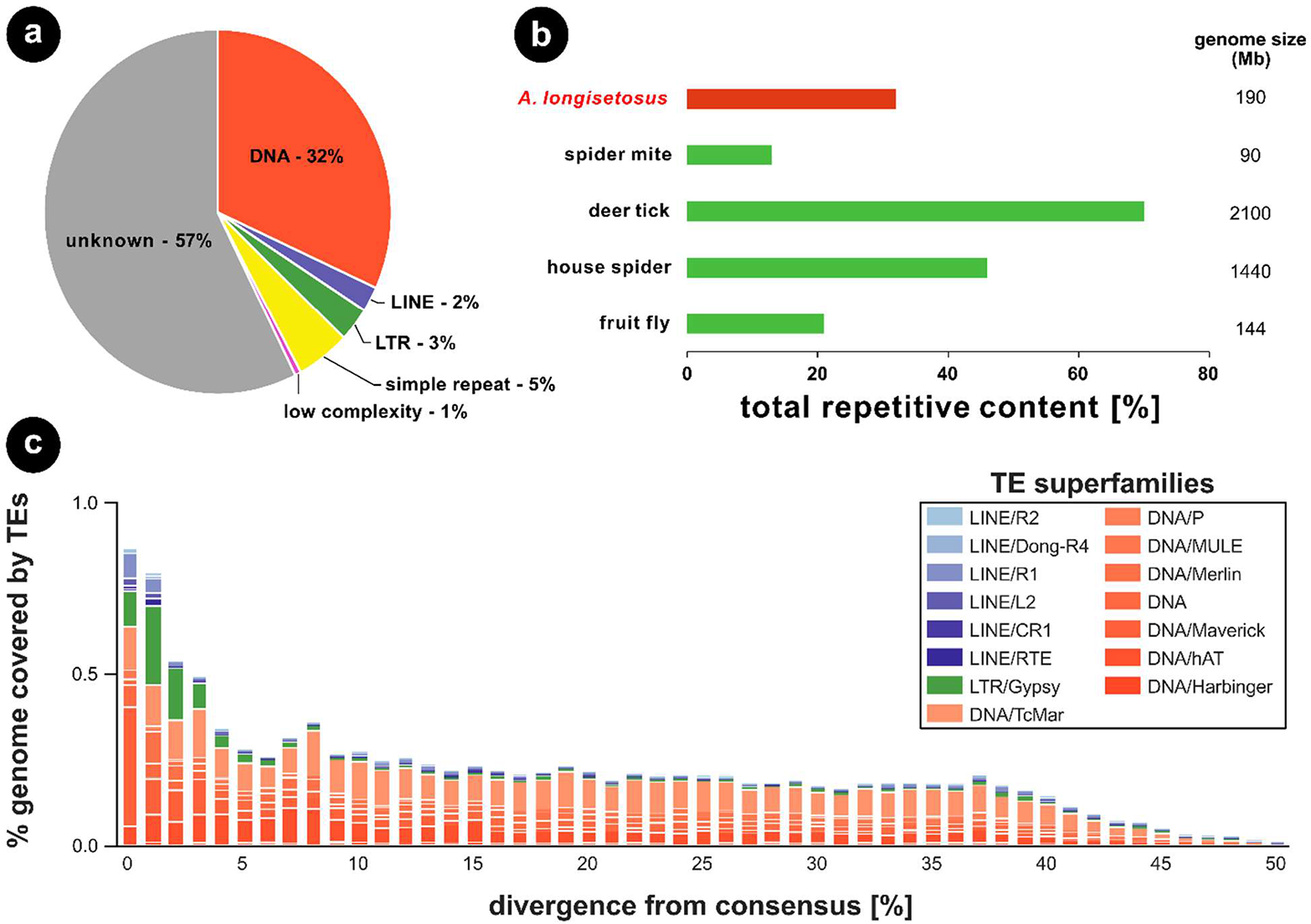
Comparison of repeat content estimations and transposable element (TE) landscape of *Archegozetes longisetosus*. a: Repetitive element categories of *Archegozetes* based on the results from RepeatModeler and MITE Tracker. LINE= long interspersed nuclear element, LTR= long terminal repeat. b: Comparison of total repetitive content among *Archegozetes*, other model chelicerates and the fly. All values are from the respective genome paper of the species, except for the fly. c: Repeat divergence plot showing TE activity through time for the major TE superfamilies of *Archegozetes*. Transposable elements with a low divergence from the consensus were recently active, while TEs diverging from the consensus depicted older activities (x-axis).

The repeat content found in other oribatid mites was lower (Bast et al. 2016), but recent studies suggest that sequencing technology, read depth and assembly quality are paramount to the capacity of identifying repeat content and TEs (Bourque et al. 2018; Panfilio et al. 2019). Hence, it is very likely the current genomic data for other Oribatida underestimates the actual total repetitive content. More low-coverage, long-read sequencing could reduce the assembly fragmentation and likely reveal a higher proportion of repeats, closer to the actual repetitiveness of oribatid genomes (Panfilio et al. 2019).

Different classes of transposable elements (TEs) are characterized by the mechanism they use to spread within genomes and are known to influence population dynamics differently (Bourque et al. 2018; Crescente et al. 2018; Finnegan 1989). We therefore analyzed the evolutionary history of TE activity in *Archegozetes* in more detail (Figure 4c). The main TE superfamilies were DNA transposons (Figure 4a and c), which seems to be a common pattern of oribatid mite genomes. For *Archegozetes*, they appear to have accumulated in the genome for a long time (i.e. they are more divergent from the consensus; (Waterston et al. 2002)) with Tc1/mariner – a superfamily of interspersed repeats DNA transposons (Bourque et al. 2018) – being the most abundant one (Figure 4c). Interestingly, we found an increase in TE activity with 0-3% sequence divergence range, indicating a recent burst (Figure 4c). This burst contained an enrichment of DNA Mavericks, which are the largest and most complex DNA transposons with homology to viral proteins (Bourque et al. 2018), but also several of retrotransposons. Among these, is the Long Terminal Repeat (LTR) gypsy retroelement (Figure 4c), which is closely related to retroviruses (Bourque et al. 2018). Like retroviruses, it encodes genes equivalent to gag, pol and env, but relatively little is known about how it inserts its DNA into the host genome (Dej et al. 1998; Havecker et al. 2004). So far, it is unknown what these TEs do in *Archegozetes*, but the recent burst in TE abundance might suggest that some changes in the genome might have happened since the became a laboratory model nearly 30 years (Heethoff et al. 2013).

### The *Archegozetes* Hox cluster

The Hox genes are a group of highly conserved transcription factor-encoding genes that are used to pattern the antero-posterior axis in bilaterian metazoans (Holland and Hogan 1988; Hrycaj and Wellik 2016). Ancestrally, arthropods likely had ten Hox genes arranged in a cluster (Hughes and Kaufman 2002). During arthropod development, the Hox genes specify the identities of the body segments, and mutations in Hox genes usually result in the transformation of segmental identities (Hughes and Kaufman 2002). The importance of Hox genes in development of metazoans makes knowledge of their duplication and disappearances important for understanding their role in the evolution of body plans (Hughes and Kaufman 2002).

Mites largely lack overt, external signs of segmentation, other than the serially arranged appendages of the prosoma (Dunlop and Lamsdell 2017). Signs of segmentation in the posterior body tagma, the opisthosoma, do exist in adult members of Endeostigmata (van der Hammen 1970). However, these segmental boundaries are largely present only in the dorsal opisthosoma, making it difficult to assess how these correspond to the ventral somites (Dunlop and Lamsdell 2017; van der Hammen 1970). Developmental genetic studies of the spider mite and *Archegozetes* suggest that acariform mites only pattern two segments in the posterior body region, the opisthosoma, during embryogenesis (Barnett and Thomas 2012; 2013b; 2018; Grbić et al. 2011). This stands in stark contrast to other studied chelicerate embryos. For example, during embryogenesis the spider Parasteatoda tepidariorum patterns twelve opisthosomal segments (Schwager et al. 2015) and the opilionid Phalangium opilio patterns seven (Sharma et al. 2012). Furthermore, a member of Parasitiformes, the tick Rhipicephalus microplus, appears to pattern eight opisthosomal segments during embryogenesis (Santos et al. 2013).

Parallel to the observation of segmental reduction in T. urticae, genomic evidence suggests that this acariform mite has lost two of its Hox genes, i.e., Hox3 and abdominal-A (*abd-A*) (Grbić et al. 2011). Interestingly, orthologs of *abd-A* in other studied arthropods pattern the posterior segments as well. A genomic comparison of arthropod Hox clusters has also shown a correlation between independent losses of *abd-A* and a reduction in posterior segmentation (Pace et al. 2016). To investigate whether the loss of segmentation in *Archegozetes* is also due to an absence in *abd-A*, we annotated its Hox cluster, paying close attention to the region between the Hox genes *Ultrabithorax (Ubx)* and *Abdominal-B (Abd-B)*, which is usually where this gene resides in other arthropods (Hughes and Kaufman 2002). Our results suggest that the *Archegozetes* Hox genes are clustered in a contiguous sequence (tig00005200_pilon, total size ∼7.5 Mbp) in the same order as suggested for the ancestral arthropod (Heethoff and Rall 2015). Furthermore, we found no sequences suggestive of an *abd-A* ortholog in *Archegozetes* (Figure 5a). These data also support the findings of a previous PCR survey that retrieved no *abd-A* ortholog in *Archegozetes* (Cook et al. 2001). Genomic evidence from the Parasitiformes Ixodes scapularis and Metaseiulus occidentalis reveal that these taxa maintain orthologs of all ten Hox genes, however in *M. occidentalis* these genes are not clustered as they are in *I. scapularis* (Gulia-Nuss et al. 2016; Hoy et al. 2016).

**Figure 5.**
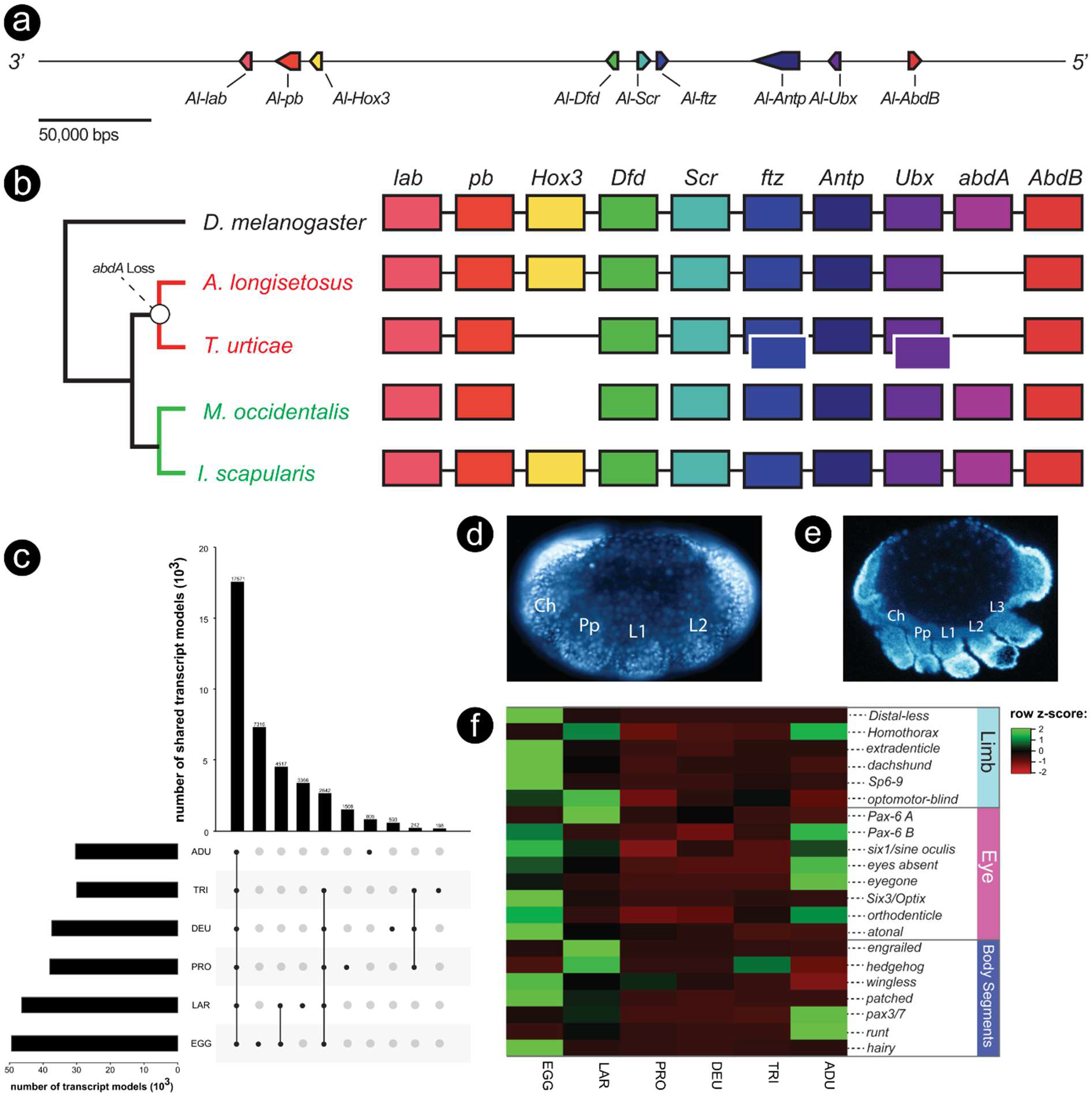
The genomic organization of the Hox genes and life-stage specific expression patters of developmental genes in *Archegozetes longisetosus*. a: Schematic of the genomic region enclosing the *Archegozetes* Hox cluster. The genomic organization of the Hox cluster is collinear, as it is in many arthropod taxa, however an abdominal-A ortholog is absent. Arrowed boxes denote the direction of transcription. The scale bar represents 50,000 base pairs. b: A comparison of the Hox cluster organization of reported members of Acari with the fruit fly *Drosophila melanogaster* as the outgroup. The last common ancestor of the parasitiform mites M. occidentalis and I. scapularis likely had an intact Hox cluster (green branches and labels), whereas abdominal-A was likely lost in the last common ancestor of acariform mites, as represented by *Archegozetes* and T. urticae (red branches and labels). Boxes with white borders represent duplicated Hox genes. Lines through the boxes indicate an intact Hox cluster. See text for further details. c: Number of transcripts shared across the different life stages of *Archegozetes*. The barplot panel on the left shows the numbers of transcripts in each stage. Exemplars of (d) early and (e) mid-germ-band embryos. Ch= chelicera; L1-3= walking legs 1-3; Pp= pedipalp. Embryos are stained with the nuclear dye DAPI and oriented with the anterior to the left of the page. f: Non-clustered heatmap showing the relative expression (row z-score based on tpm) patterns of putative limb, eye, and body segmentation genes throughout the embryonic, larval instars, and adult stages of *Archegozetes*. **See supplementary Table S3** for average tpm values. Life stages (for c and f): EGG= egg; LAR= larva; PRO= protonymph; DEU= deutonymph; TRI= tritonymph; ADU= adult.

Taken together, these observations suggest that the last common ancestor of acariform mites likely lost its abdominal-A gene as well as experiencing a reduction in opisthosomal segmentation (Figure 5b). Alternatively, these shared losses of *abd-A* may be due to convergence due to similar selective pressures favoring a reduction in body size. The dorsal, external segmentation of endeostigmatid mites does not necessarily contradict the hypothesis of a loss of *abd-A* at the base of the acariform mites. As Hox genes are usually deployed after the genetic establishment of segments in arthropods (Hughes and Kaufman 2002), the opisthosomal segments in endeostigmatid mites may still develop in the absence of *abd-A*. However, this hypothesis needs further testing with observations of segmental gene expression in endeostigmatids as well as additional acariform species.

### Life-stage specific RNA expression patterns

Developmental and gene expression data from *Archegozetes* embryos (Figure 5 d and e) have elucidated many of the potential mechanisms driving the morphogenesis of manydevelopmental peculiarities. These peculiarities include the suppression of the fourth pair of walking legs during embryogenesis as well as the reduction of opisthosomal segmentation (Barnett and Thomas 2012; 2013a; 2013b; 2018; Telford and Thomas 1998; Thomas 2002). In typical acariform mites, embryogenesis ends with the first instar, the prelarva, which usually remains within the egg chorion, as in Archegozeetes. Hatching releases the second instar, the larva, which is followed by three nymphal instars (proto-, deutero- and tritonymph) and the adult, for a total of six instars. (Heethoff et al. 2007). Thus far, methodological limitations have made it difficult to examine how mite segmentation and limb development progress throughout these instars.

To this end, we used RNAseq to calculate the transcripts per million (tpm) values of genes known to be, or suspected to be, involved in limb development and segmentation throughout the six different instars of *Archegozetes*. Prior to comparing these tpm values, gene orthology was confirmed via phylogenetic analyses (supplementary Figures S4-S11; see Table S2 for phylogenetic statistics and Table S3 for tpm values). Regarding the total number of genes expressed across the different life stages, we found that earlier instars generally expressed a higher number of genes (Figure 5c). While most expressed genes were shared across all instars, more transcripts were shared between the eggs and the larvae and among all five juvenile instars. Additionally, we found that earlier instars expressed a larger number of stage-specific genes as compared to later instars and adults (Figure 5c).

Gene expression, SEM and time-lapse data have revealed that the development of the fourth pair of walking legs in *Archegozetes* is suppressed until after the larval instar (Barnett and Thomas 2012; 2018; Telford and Thomas 1998). The resulting larva is thus hexapodal (see also embryo in Figure 5e), which constitutes a putative synapomorphy of Acari, if they are monophyletic (Dunlop and Alberti 2008). In arthropods, the development of the limbs is generally accomplished via the activity of highly conserved regulatory genes, termed the “limb gap genes.” These genes are expressed along their proximo-distal axes to establish the specific identities of the limb podomeres. The limb gap genes include extradenticle (exd) and homothorax (hth), which act together to specify the proximal limb podomeres, dachshund (dac), which specifies the medial podomeres, and Distal-less (Dll) which specifies the distal-most podomeres. It was previously shown that the deployment of these genes in the anterior appendages of *Archegozetes*, i.e., the chelicerae, pedipalps and first three pairs of walking legs (Figure 5d and e), is similar to that of other chelicerate taxa (Barnett and Thomas 2013a; Schwager et al. 2015; Sharma et al. 2015). However, in the anlagen of the fourth pair of walking legs, only the proximal-specifying genes, exd and hth, are expressed (Barnett and Thomas 2013a).

Whether the limb gap genes are re-deployed during the transition from the prelarval to larval instars in order to activate the development of the fourth pair of walking legs remains an open question. We therefore compared the average tpm values of verified limb gap genes (i.e., Al-Dll, Al-Hth, Al-exd, and Al-dac (Barnett and Thomas 2013a)) in embryos and at each instar stage (Figure 5f). We also compared the tpm values of the *Archegozetes* orthologs of Sp6-9 and optomotor blind, genes shown to be involved in limb formation in spiders (Heingård et al. 2019; Königsmann et al. 2017). We hypothesized that limb development genes would show high expression in the larval stage leading to the development of the octopodal protonymph. We did observe an increase in the tpm averages of Al-hth as well as Al-optomotor-blind, however the aforementioned limb gap gene expression levels were similar between these instars (Figure 5f). Taken together, these genes may not be up-regulated for the formation of the fourth pair of walking legs between these two instars.

Chelicerate embryos segment their bodies through a “short/intermediate germ” mechanism, whereby the anterior (prosomal) segments are specified asynchronously (Schwager et al. 2015). This usually occurs well before the sequential addition of posterior segments from a posterior growth zone. Based on neontological and paleontological data, chelicerate arthropods may have ancestrally had an opisthosoma comprised of 12 or more segments (Dunlop and Selden 1998; Dunlop 2010; Dunlop and Lamsdell 2017). Embryonic expression data for the segment polarity genes, those genes that delineate the boundaries of the final body segments, have shown that in most studied chelicerate embryos opisthosomal segments are delineated during embryogenesis (Dunlop and Lamsdell 2017; Schwager et al. 2015). However, as discussed above, expression data in *Archegozetes* embryos suggest that only two opisthosomal segments are patterned during embryogenesis (Barnett and Thomas 2012; 2018); this indicates that mites have significantly reduced their number of opisthosomal segments either by loss or by fusion. Further complicating this is the observation that many mites add segments as they progress through the larval instars, a phenomenon known as anamorphic growth (Dunlop and Lamsdell 2017).

To determine by what genetic process *Archegozetes* may add segments during post-embryonic ontogeny, we assessed the expression of known chelicerate and arthropod segmentation genes in each instar transcriptome (Figure 5f) (Schwager et al. 2015). We observed an up-regulation of the segmentation genes hedgehog and engrailed in the larvae, as well as the slight up-regulation of patched and pax3/7. Furthermore, the segmentation gene wingless was slightly up-regulated in the protonymph, as well as a slight up-regulation of hedgehog in the tritonymph. Lastly, we found that transcripts of the genes pax3/7 and runt were up-regulated in adults. These results suggest that *Archegozetes* does pattern body segments during the progression through the it’s instars similar to other Chelicerata.

Another peculiarity of *Archegozetes* is that these mites lack eyes (see more details below). Eye loss has been documented in other arachnid clades, including independently in other members of Acari (Evans 1992; Walter and Proctor 1999), and it has been recently demonstrated that a species of whip spider has reduced its eyes by reducing the expression of retinal determination genes that are shared throughout arthropods (Gainett et al. 2020). We sought to determine if eye loss in *Archegozetes* also is associated with the reduced expression of these genes (see also analysis of photoreceptor genes below). The genes, which have been shown to be expressed in the developing eyes of spiders and whip scorpions, include Pax-6, six1/sine oculis (so), eyes absent (eya), Eyegone, Six3/Optix, and atonal (Gainett et al. 2020; Samadi et al. 2015; Schomburg et al. 2015). We also followed the expression of Al-orthodenticle, a gene previously shown to be expressed in the ocular segment of *Archegozetes* (Telford and Thomas 1998). Surprisingly, all of these genes, excluding the Pax-6 isoform A and eyegone, are indeed expressed during embryogenesis (Figure 5f). Aside from the larval expression of the Pax-6 isoform A during the larval stage, these eye-development genes remain quiescent until the adult stage, where all but Pax-6 isoform A, six3 and atonal are up-regulated (Figure 5f). These results are exceedingly surprising, given the conserved role of genes in retinal patterning. They suggest a novel role for these genes, or alternatively, these expression patterns could be the result of early expression of a retinal determination pathway followed by negative regulation by other genes to suppress eye development.

### Photoreceptor and chemosensory system of *Archegozetes longisetosus*

Unlike insects and crustaceans, chelicerates do not have compounds eyes – with horseshoe crab being an exception. Generally, mites are eyeless or possess one or two pairs of simple ocelli (Alberti and Coons 1999; Alberti and Moreno-Twose 2012; Exner 1989; Harzsch et al. 2006; Patten 1887). Ocelli are common in Prostigmata and Endeostigmata, among Acariformes, as well Opilioacarida – the most likely sister group to the Parasitiformes – but are absent in most Oribatida, Astigmata, Mesostigmata and ticks (Norton and Franklin 2018; Norton and Fuangarworn 2015; Walter and Proctor 1998; Walter and Proctor 1999). This suggests that the presence of eyes might be an ancestral condition for both Acariformes and Parasitiformes, while more derived mites rely largely on chemical communication systems (Alberti and Coons 1999).

In oribatid mites, detailed morphological and ultrastructural investigations have suggested that setiform sensilla are the most obvious sensory structures (Figure 6a) (Alberti 1998; Alberti and Coons 1999; Walter and Proctor 1999). The trichobothria are very complex, highly modified (e.g. filiform, ciliate, pectinate, variously thickened or clubbed) no-pore setae which are anchored in a cup-like base and likely serve as mechanosensory structures. In contrast, the setal shafts of solenidia and eupathidia (Figure 6a) both possess pores (Alberti 1998; Alberti and Coons 1999; Walter and Proctor 1999). Solenidia have transverse rows of small pores visible under a light microscope and likely function in olfaction, while the eupathidia have one or several terminal pores and likely are used as contact/gustatory sensilla (Figure 6a) (Alberti 1998; Alberti and Coons 1999). Previous work demonstrated that oribatid mites indeed use olfactory signals in the context of chemical communication and food selection (Brückner et al. 2018a; Brückner et al. 2018b; Heethoff et al. 2011a; Heethoff and Raspotnig 2012; Raspotnig 2006; Shimano et al. 2002).

**Figure 6.**
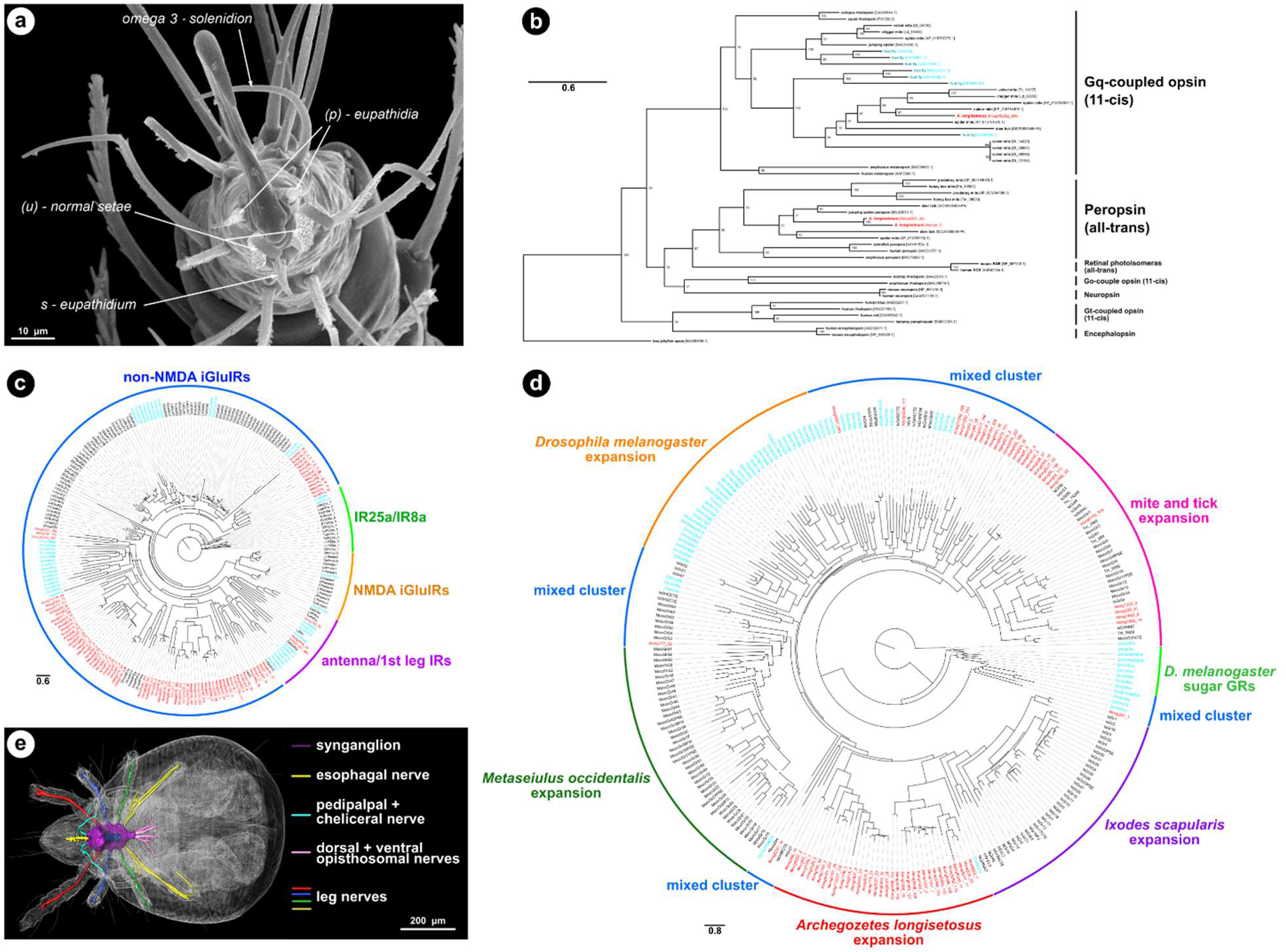
The sensory systems of *Archegozetes longisetosus* and phylogenetic analysis of selected photoreceptor and chemosensory genes. **a**: Scanning electron micrograph (SEM) showing the end of tarsus on *Archegozetes*’ first leg. Images shows normal setae, but also modified chemosensory setae, namely eupathidia, both paired (p) and single (s), as well as an omega-3 solenidium. SEM picture courtesy of Michael Heethoff. **b**: Phylogeny and classification of opsin genes across the Metazoa, including those of several Chelicerata. The tree was constructed using a maximum likelihood approach and rooted with a jelly fish opsin. *Archegozetes* sequences are depicted in red, Drosophila in turquoise; branch length unit is substitutions per site. **c**: Maximum likelihood phylogeny of ionotropic receptors and ionotropic glutamate receptors of *Archegozetes* (Along), *Dinothrombium* (Dt), *Leptothrombidium* (Ld), *Tetranychus* (Tu) and *Drosophila* (Dmel). IR25a/IR8a and antenna/1^st^ leg IRs contain genes with known chemosensory function in Drosophila. The tree was rooted to the middle point; *Archegozetes* sequences are depicted in red, Drosophila in turquoise; branch length unit is substitutions per site. Bootstrap values can be found in the supplementary Figure S13. **d**: Maximum likelihood phylogenetic tree of gustatory receptors of *Archegozetes* (Along), Ixodes (Is), Tropilaelaps (Tm), Metaseiulus (Mocc) and Drosophila (Dmel). The tree was rooted to the middle point; *Archegozetes* sequences are depicted in red, Drosophila in turquoise; branch length unit is substitutions per site. Bootstrap values can be found in the supplementary Figure S14. **e**: Combined image of volume rendering (grey) and reconstructed nervous system of *Archegozetes* in dorsal view. Color-code corresponds to different parts of the nervous system, as depicted in the legend. The blue structure in the middle of the synganglion is the part of the esophagus which penetrates the synganglion. Scale bar: 200 µm. Image courtesy of Sebastian Schmelzle based on data in (Hartmann et al. 2016).

Interestingly, detailed morphological and ultrastructural studies showed that light-sensitive organs exist in some Palaeosomata and Enarthronota (probably true eyes) as well as in Brachypylina (the secondary lenticulus), representing lower and highly derived oribatid mites, respectively (Alberti and Coons 1999; Alberti and Moreno-Twose 2012; Norton and Franklin 2018; Norton and Fuangarworn 2015). *Archegozetes* and most other oribatids, however, are eyeless, yet there is scattered experimental and some anecdotal evidence that even these mites show some response to light and seem to avoid it (‘negative phototropism’ or ‘negative phototaxis’) (Madge 1965; Trägårdh 1933; Walter and Proctor 1999; Woodring 1966). Hence, we mined the genome of *Archegozetes* for potential photoreceptor genes and found two genes of the all-trans retinal peropsin class and one gene related to spider mite rhodopsin-7-like gene (Figure 6b). Peropsin-like genes are also present in other eyeless ticks. In jumping spiders it encodes for nonvisual, photosensitive pigments, while rhodopsin-7 may be involved in basic insect circadian photoreception (Eriksson et al. 2013; Koyanagi et al. 2008; Nagata et al. 2010; Senthilan et al. 2019; Senthilan and Helfrich-Förster 2016; Shen et al. 2011). Taken together, this might suggest that eyeless species like *Archegozetes* use peropsin- and rhodopsin-7-like genes for reproductive and diapause behaviors, or to maintain their circadian rhythm, as well as negative phototaxis.

However, the main sensory modality soil mites use is chemical communication via olfaction (Alberti 1998; Alberti and Coons 1999; Brückner et al. 2018a; Brückner et al. 2018b; Raspotnig 2006; Shen et al. 2011; Walter and Proctor 1999). In contrast to insects, but similar to crustaceans and Myriapoda, mites do not have the full repertoire of chemosensory classes, they are missing odorant receptors and odorant-binding proteins (Table 2) (Dong et al. 2017; Dong et al. 2018; Hoy et al. 2016; Maraun et al. 2007; Raspotnig 2009; Sánchez-Gracia et al. 2009; Sánchez-Gracia et al. 2011; Vieira and Rozas 2011). Although chemosensory protein (CSP) encoding genes are absent in most mite genomes, we identified one gene encoding for such a protein in *Archegozetes* and one CSP has been previously found in the deer tick (Table 2). Hence, *Archegozetes* should primarily rely on gustatory receptors (GRs) and ionotropic receptors (IRs). Both the number of GRs (68 genes; Figure 6d) and IRs (3 genes; Figure 6c) was very well within the range of most mites and ticks and there was no evidence for any massive chemoreceptor expansion like in the spider mite (Table 2) (Ngoc et al. 2016). This was surprising because *Archegozetes*, like other acariform mites have many multiporous solenidia, present on all legs and the palp, but appear to only have a limited number of chemoreceptors.

**Table 2.** Comparison of chemosensory receptor repertoires between *Archegozetes longisetosus* and other arthropods. GR= gustatory receptor, OR= odorant receptor, IR= ionotropic receptor, OBP= odorant binding protein, CSP= chemosensory protein.

|  | **Chemosensory receptors** |  |  |  |  |
| --- | --- | --- | --- | --- | --- |
|  | **GR** | **OR** | **IR** | **OBP** | **CSP** |
| *A. longesitosus* | 68 | 0 | 3 | 0 | 1 |
| spider mite | 689 | 0 | 4 | 0 | 0 |
| deer tick | 60 | 0 | 22 | 0 | 1 |
| house spider | 634 | 0 | 108 | 4 | 0 |
| fruit fly | 73 | 62 | 66 | 51 | 4 |

Canonical ionotropic glutamate receptors (iGluRs) are glutamate-gated ion channels with no direct role in chemosensation, which come in two major subtypes: either NMDA iGluRs which are sensitive to N-methyl-D-aspartic acid (NMDA) or non-NMDA iGluRs. The latter group – at least in Drosophila – seems to have essential functions in synaptic transmission in the nervous system and have been associated with sleep and vision (Benton et al. 2009; Croset et al. 2010; Ngoc et al. 2016; Sánchez-Gracia et al. 2009; Sánchez-Gracia et al. 2011). None of the IRs we found in the *Archegozetes* genome belonged to the NMDA iGluRs and most were classified as non-NMDA iGluRs (Figure 6c). Nothing is known about their functions in mites. It is, however, likely that they perform similar tasks in synaptic transmission in the brain and musculature. In Drosophila a specific set of chemosensory IRs, which do not bind glutamate, respond to acids and amines (IR25a), but also to temperature (IR21a, IR93a). For *Archegozetes* we found 3 IRs, like IR21a and IR93a of Drosophila, which fell into the antenna/1st leg IRs category (Table 2; Figure 6c) (Budelli et al. 2019; Knecht et al. 2016; Rytz et al. 2013). This is consistent with an assumed limited contribution of IRs to the perception of chemical cues. Furthermore, it is so far unclear whether these IRs are expressed in the first pair of legs (Figure 6a and c) in *Archegozetes*, but similar genes seem to be expressed in the legs of other mite species, which could suggest a similar function as in the fruit fly.

GRs are multifunctional proteins and at least in insects they are responsible for the perception of taste, heat or volatile molecules (Montell 2009). In *Archegozetes* we found 68 GRs, over half of which belonged to a species-specific expansion of the GR gene family (Figure 6d). Generally, it is unclear if GRs in *Archegozetes* and other mites have similar functions as in insects, but the GR gene family is heavily expanded in many acariform mites and also is present in ticks (Table 2), suggesting an important biological role (Barrero et al. 2017; Dong et al. 2017; Dong et al. 2018; Gulia-Nuss et al. 2016; Hoy et al. 2016; Ngoc et al. 2016). This is supported by experimental evidence which suggested that ticks and other mites, including *Archegozetes*, use chemical cues to find their host, communicate or discriminate food (Barrero et al. 2017; Brückner et al. 2018a; Brückner et al. 2018b; Bunnell et al. 2011; Gulia-Nuss et al. 2016; Kuwahara 2004; Raspotnig 2006; Yunker et al. 1992).

In general, not much is known about the nervous and sensory system of oribatid mites, or about sensory integration or the neuronal bases of their behavior (Alberti 1998; Alberti and Coons 1999; Norton 2007). Modern methods like Synchrotron X-ray microtomography (SRμCT) recently made it possible to investigate the organization and development of the nervous systems of oribatid mites (Figure 6e; (Hartmann et al. 2016)) and here we provide the first genomic resource for the investigation of the photo- and chemosensory systems of Oribatida (Figure 6b-d).

### Horizontal gene transfer event sheds light on oribatid feeding biology

Horizontal gene transfer (HGT) is common among mites and other soil organisms (Dong et al. 2018; Faddeeva-Vakhrusheva et al. 2016; Grbić et al. 2011; Mayer et al. 2011; Wu et al. 2017; Wybouw et al. 2018). In some cases, genes that had been horizontally transferred now have pivotal biological functions. For instance, terpene and carotenoid biosynthesis genes in trombidiid and tetranychid mites, respectively, are found nowhere else in the animal kingdom (Altincicek et al. 2012; Dong et al. 2018). Yet they show high homology with bacterial (terpene synthase) or fungal (carotenoid cyclase/synthase/desaturase) genes, suggesting horizontal gene transfer from microbial donors (Altincicek et al. 2012; Dong et al. 2018). At least the carotenoid biosynthesis genes in spider mites still code for functional enzymes and equip these phytophages with the ability to de novo synthesize carotenoids, which can induce diapause in these animals (Altincicek et al. 2012). Soil microarthropods like collembolans show numbers of horizontally transferred genes that are among the highest found in Metazoan genomes, exceeded only by nematodes living in decaying organic matter (Crisp et al. 2015; Faddeeva-Vakhrusheva et al. 2016; Wu et al. 2017). Interestingly, many HGT genes found in collembolans are involved in carbohydrate metabolism and were especially enriched for enzyme families like glycoside hydrolases, carbohydrate esterases or glycosyltransferases (Faddeeva-Vakhrusheva et al. 2016; Wu et al. 2017). All three enzyme families are involved in the degradation of plant and fungal cell walls (Gilbert 2010; Latgé 2007). Hence, it has been hypothesized that cell-wall degrading enzymes acquired by HGT are beneficial for soil organisms as it allowed such animals to access important food source in a habitat that is highly biased towards polysaccharide-rich resources (Faddeeva-Vakhrusheva et al. 2016; Faddeeva-Vakhrusheva et al. 2017; Mitreva et al. 2009; Wu et al. 2017).

To assess the degree of HGT in *Archegozetes* we first used blobtools (v1.0) (Laetsch and Blaxter 2017) to generate a GC proportion vs read coverage plot of our genome assembly, in order to remove scaffolds of bacterial origin (Figure 7a; 438 contigs of ∼ 8.5 Mb of contamination). Of the remaining scaffolds, candidate HGTs were identified using the Alien Index (Flot et al. 2013; Thorpe et al. 2018), where HGTs are those genes with blast homology (bit score) closer to non-metazoan than metazoan sequences (supplementary Table S4). We further filtered these HGT candidates to remove those that overlapped predicted repeats by ≥ 50%, resulting in 617 genes. As HGT become integrated into the host genome, they begin to mirror features of the host genome, including changes in GC content and introduction of introns (Lawrence 1997). Comparing the GC content of the HGT candidates showed two distinct peaks, one at 54.3% and the other at 35.2%, slightly higher than the remaining *Archegozetes* genes, GC content of 31.5% (Figure 7b). Of the 407 HGT genes that shared similar GC content to the host genome, 73.5% had at least one intron (Table S4). In a final step, we used the gene expression data (RNAseq) to filter the list of all putative HGT genes and only retained candidates that were expressed in any life stage of *Archegozetes* (n= 298 HGT genes).

**Figure 7.**
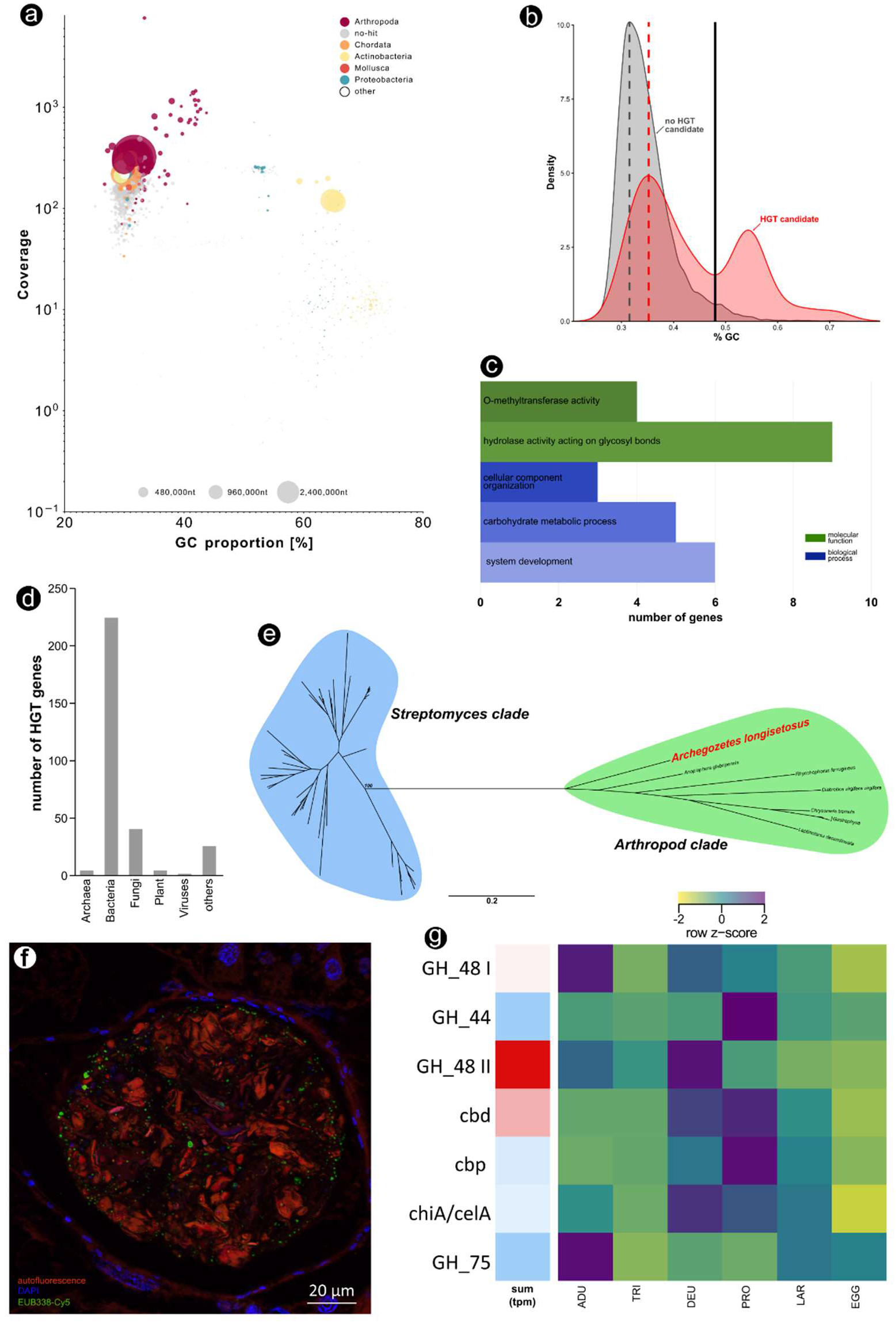
Horizontal gene transfer (HGT) and implications for the feeding biology of *Archegozetes longisetosus*. a: Blob-plot of the long-read genome assembly contigs plotting the read coverage against GC proportion [%]. Contigs are colored according to the taxonomic order of their best Megablast hit to the NCBI nucleotide database. Size of circle corresponds to the nucleotides per contigs. b: Comparison of the GC content of HGT and non-HGT genes. HGT genes shifted towards the host genome GC content indicate integration within the host genome while the higher GC content HGT genes might be the product of relatively recent HGT events. c: Enrichment of functional categories (GO terms) describing the molecular functions and biological processes related to the HGT candidate genes. d: Taxonomic origin of HGT. The category “others” includes mostly protozoan donor genes among other Eukaryotes. e: Unrooted maximum-likelihood tree of glycoside hydrolase family 48 members (GH_48) from Streptomyces bacteria and HGT genes from other arthropods as well as *Archegozetes* (GH_48 II). Bootstrap values and the full tree can be found in the supplementary Figure S15. The scale bar denotes substitutions per site. f: Fluorescence in *situ* hybridization (FISH) micrograph of a food bolus in the mites’ alimentary tract. The food material (wheat grass power) is enclosed in a peritrophic membrane and there is a high bacterial prevalence in the food bolus. Image courtesy of Benjamin Weiss and Martin Kaltenpoth. g: RNAseq support of HGT candidates related to cell wall degrading enzymes. The first block (single column) shows the overall RNA expression (tpm) of the HGT in all life stages; red denotes high total expression, while blue depicts low total expression. The second block (six columns) shows the expression (row z-score based on tpm) of the same HGT candidates across the different life stages of *Archegozetes*. Abbreviations: GH_48= glycoside hydrolase family 48, GH_44= glycoside hydrolase family 44, cbd= cellulose-binding domain, cbp= cellulose-binding protein, chiA/celA = chitinase/cellulase, GH_75= glycoside hydrolase family 75.

The majority of HGT candidates were of bacterial origin (75.2%), followed by genes likely acquired from fungi (13.4%), while transfer from Archaea, plants, virus and other sources was comparatively low (Figure 7d). This composition of HGT taxonomic origin is different from genes found in collembolans, which appear to have acquired more genes of fungal and protist origin (Faddeeva-Vakhrusheva et al. 2016; Faddeeva-Vakhrusheva et al. 2017; Wu et al. 2017). Subsequently, we performed an over-representation analysis of GO terms associated with these genes. We found an over-representation of genes with GO terms related to methyl transfer reactions and breaking glycosidic bonds (molecular function; Figure 7c) as well as carbohydrate metabolism, among others (biological process; Figure 7c) providing a first line of evidence that *Archegozetes* possess HGT related to plant- and fungal cell wall degradation similar to springtails.

As mentioned previously, oribatid mites are among the few Chelicerata that ingest solid food and are primary- and secondary decomposers feeding on dead plant material and fungi (Cohen 1995; Dunlop and Alberti 2008; Heethoff and Norton 2009; Maraun et al. 2011; Norton 2007; Shultz 2007). It was argued for decades that the enzymes necessary to break down these polysaccharide-rich resources originate from the mite’s gut microbes (Siepel and de Ruiter-Dijkman 1993; Smrž 1992; Smrž 2000; Smrž and Čatská 2010; Smrž and Norton 2004; Stefaniak 1976; 1981). Microbes might be mixed with the food in the ventriculus and digest it while passing through the alimentary tract as food boli enclosed in a peritrophic membrane (see Figure 7f for an example) (Stefaniak 1976; 1981). However, screening the HGT candidate list for potential cell-wall degrading enzymes and mapping their overall and life-stage specific expression in *Archegozetes* using the RNAseq reads, revealed at least seven HGT genes related to polysaccharide breakdown (Figure 7g). We found that specifically members of the glycoside hydrolases family 48 and cellulose-binding domain genes showed high expression in most life stages - the egg being an obvious exception (Figure 7g). Moreover, the majority of these genes were flanked by a predicted metazoan gene, suggesting host transcriptional regulation (Table S4).

In a last step we blasted the highly expressed HGT candidates (Figure 7g) against the non-redundant protein sequence database, aligned the sequences with the highest alignment score. Eventually, we performed a phylogenetic maximum likelihood analysis. For the highest expressed HGT related to cell-wall-degrading enzymes (glycoside hydrolases family 48 gene II), we recovered that the *Archegozetes* sequences was well nested within a clade of GH 48 sequences from herbivores beetles (McKenna et al. 2019), which appear to be related to similar genes from various Streptomyces (Figure 7e) and we reconstructed similar phylogenies for other highly expressed HGT candidates (supplementary Figure S12). All the sequences of beetle glycoside hydrolases family 48 members (Figure 7e) were included in recent studies arguing for a convergent horizontal transfer of bacterial and fungal genes that enabled the digestion of lignocellulose from plant cell walls in herbivores beetles (McKenna et al. 2016; McKenna et al. 2019). They showed that phytophagous beetles likely acquired all genes of the GH 48 family from Actinobacteria (including Streptomyces) (McKenna et al. 2019) and our phylogenetic analysis (Figure 7e) revealed the same pattern as well as a highly similar tree topology (compare to Fig 3B in (McKenna et al. 2019)).

Overall, our findings indicate that genes encoding for enzymes in *Archegozetes* capable of degrading plant and fungal cell walls were likely horizontally transferred from bacteria (likely Streptomyces). Bacterial symbionts and commensal living in the mites’ gut are still likely to contribute to the breakdown of food (Figure 7f). Yet, the high expression of genes encoding cell-wall degrading enzymes (Figure 7g) as well as the evolutionary analyses of such genes (Figure 7e) suggest that *Archegozetes* – and potentially many other oribatid mites – are able to exploit polysaccharide-rich resources like dead plant material or chitinous fungi without microbial aid. Enzymological and microscopical investigation of *Archegozetes* have suggested that certain digestive enzymes (chitinase and cellulase) are only active when the mites consume a particular type of food (e.g. algae, fungi or filter paper) (Smrž and Norton 2004). These results were interpreted as evidence that these enzymes are directly derived from the consumed food source (Smrž and Norton 2004). By contrast, we argue that this instead confirms our findings of HGT: upon consumption of food containing either chitin or cellulose, gene expression of polysaccharide-degrading enzymes starts, and proteins can readily be detected. Further enzymological studies have placed oribatid mites in feeding guilds based on carbohydrase activity and also found highly similar enzyme activity between samples of mites from different times and locations (Luxton 1972; 1979; 1981; 1982; Siepel and de Ruiter-Dijkman 1993). Future functional studies can disentangle the contribution of the host and microbes to cell wall digestion and novel metabolic roles of the HGTs identified here.

### Biosynthesis of monoterpenes – a common chemical defense compound class across oribatid mite

Oribatid and astigmatid mites are characterized by a highly diverse spectrum of natural compounds that are produced by and stored in so-called oil glands (for an example see Figure 8a) (Heethoff et al. 2016; Raspotnig 2009; Raspotnig et al. 2011). These paired glands are located in the opisthosoma (i.e. the posterior part of chelicerate arthropods, analogous to the abdomen of insects) and are composed of a single-cell layer invagination of the cuticle (Figure 8f). As previously mentioned, mites use chemicals produced by these glands to protect themselves against environmental antagonists (predators or microbes) or use them as pheromones (Brückner et al. 2015; Heethoff et al. 2011a; Heethoff and Rall 2015; Heethoff and Raspotnig 2012; Raspotnig 2006; 2009; Shimano et al. 2002). The monoterpene aldehyde citral – a stereoisomeric mixture of geranial ((E)-3,7-dimethylocta-2,6-dienal) and neral ((Z)-3,7-dimethylocta-2,6-dienal) – and its derivatives are widely detected compounds in glandular secretions of oribatids and astigmatids (Koller et al. 2012; Kuwahara 2004; Kuwahara et al. 2001; Raspotnig et al. 2004; Sakata 1997; Sakata and Norton 2001; Sakata and Norton 2003; Sakata et al. 1995). These monoterpenes have been called “astigmatid compounds” (Sakata and Norton 2001) as they characterize the biochemical evolutionary lineage of major oribatid mite taxa (Mixonomata and Desmonomata) and almost all investigated astigmatid mites (Alberti 1984; Kuwahara 2004; Raspotnig 2009; Sakata 1997; Sakata and Norton 2001).

**Figure 8.**
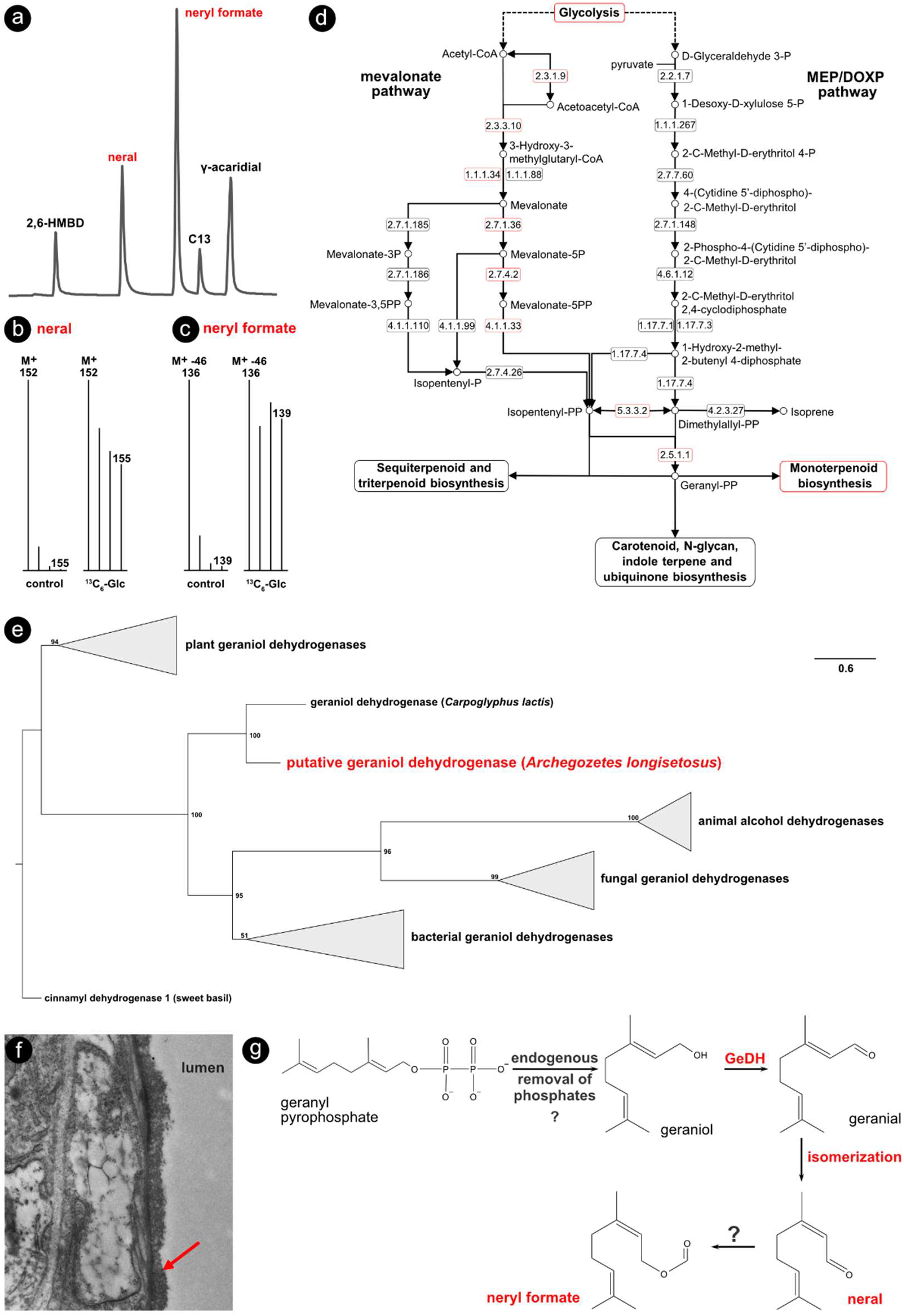
Reconstruction of the biosynthetic pathway leading to monoterpenes in *Archegozetes* longisetosus. a: Representative gas chromatogram of the mite’ gland content; in order of retention time: 2-hydroxy-6-methyl-benzaldehyde (2,6-HMBD), neral ((Z)-3,7-dimethylocta-2,6-dienal) neryl formate ((Z)-3,7-dimethyl-2,6-octadienyl formate), tridecane, 3-hydroxybenzene-1,2-dicarbaldehyde (γ-acaridial). Further alkanes/alkenes (pentadec-7-ene, pentadecane, heptadeca-6,9-diene, heptadec-8-ene, heptadecane) are not shown. Monoterpenes are marked in red. b and c: Representative mass spectra of neral (b) and neryl formate (c) extracted from defensive glands of mites fed with unlabeled wheatgrass powder (control), or wheatgrass infused with ^13^C_6_-labelled glucose recorded in single-ion mode. The mass spectra for neral (b) shows the M+-ion series, while the spectra for neryl formate (c) show the diagnostic ion series at [M-46]^+^. Mites fed with the ^13^C_6_ glucose infused wheatgrass showed enriched ions. d: KEGG reference pathway map for terpenoid backbone biosynthesis. Mapping genes from the *Archegozetes* genome encoding for pathway enzymes (labeled in red) revealed that the mite can produce geranyl pyrophosphate (GPP) via the mevalonate pathway from precursors provided by glycolysis. Enzymes names correspond to EC numbers: 2.3.1.9= acetyl-CoA C-acetyltransferase; 2.3.3.10= hydroxymethylglutaryl-CoA synthase; 1.1.1.34= hydroxymethylglutaryl-CoA reductase; 2.7.1.36= mevalonate kinase; 2.7.4.2= phosphomevalonate kinase; 4.1.1.33= diphosphomevalonate decarboxylase; 5.3.3.2= isopentenyl-diphosphate delta-isomerase; 2.5.1.1= farnesyl diphosphate synthase. e: Maximum-likelihood tree based on an alignment of plant, fungal and bacterial geraniol dehydrogenases, animal alcohol dehydrogenase and two mite (*Carpoglyphus lactis* and *Archegozetes*) geraniol dehydrogenases (GeDH). Bootstrap values (based on 1000 replicates) are indicated along branches and the scale bar denotes substitutions per site. The tree was rooted by the outgroup cinnamyl dehydrogenase from sweet basil. f: Ultrastructure of the gland-tissue of *Archegozetes*, as observed by transmission electron microscopy (TEM). Red error shows the border between the gland cell and the glandular lumen. TEM picture courtesy of Michael Heethoff. g: Proposed biochemical pathway scenario leading to neral and neryl formate in *Archegozetes* starting with GGP from the terpenoid backbone biosynthesis.

The chemical cocktail released by *Archegozetes* consists of a blend of 10 compounds (Figure 8a) including two terpenes (approx. 45%)– neral and neryl formate – six hydrocarbons (approx. 15%) and two aromatic compounds (approx. 40%) (Brückner and Heethoff 2017; Sakata and Norton 2003). The hydrocarbons likely serve as solvents, while the terpenes and aromatics are bioactive compounds used in chemical alarm and defense (Heethoff et al. 2011a; Raspotnig 2006; Sakata and Norton 2003; Shimano et al. 2002). Recently, it was shown that *Archegozetes* synthesizes the two aromatic compounds using a polyketide-like head-to-tail condensation of (poly)-β-carbonyls via a horizontally acquired putative polyketide synthetase (Brückner et al. 2020). Studies in Astigmata found that the monoterpenes of these mites appeared to be made de novo from (poly)-β-carbonyls as well and one study identified a novel geraniol dehydrogenases (GeDH), unrelated to those of bacteria, in Carpoglyphus lactis (Morita et al. 2004; Noge et al. 2005; Noge et al. 2008). To learn about the biosynthesis of astigmatid compounds in *Archegozetes* and demonstrate the mite’s applicability as research model for biochemical pathway evolution, we used the novel genomic resources presented in this study.

First, we delineated the basic biochemical reactions likely to happen in the *Archegozetes* gland through a stable-isotope labeling experiment. We supplemented the diet of the mite with food containing 25% heavy ^13^C_6_ D-glucose and 10% antibiotics (a combination of three different antibiotics was fed, because this mixture is able to eliminate nearly all qPCR and FISH detectable bacteria found on the food and in the alimentary tract (Brückner et al. 2020)). To examine the incorporation of heavy ^13^C_6_ D-glucose and its metabolic products into neral (Figure 8b) and neryl formate (Figure 8c), we compared selected fragment ions (M^+^ and M^+^-46, respectively) using single ion mass spectrometry. Both neral and neryl formate showed consistent enrichment in their M^+^ to [M+4]^+^ and [M-46]^+^ to [M-46+4]^+^-ion series, indicating that *Archegozetes* used glycolysis breakdown products of ^13^C_6_ D-glucose for the biosynthesis of their monoterpenes. We then used the OGS mapped to KEGG metabolic pathways (Kanehisa et al. 2007) to reconstruct the backbone synthesis of terpenes in *Archegozetes* (Figure 8d). We found mapped mite genes, which suggest that *Archegozetes* synthesizes geranyl pyrophosphate (GPP) – the input substrate for further monoterpene synthesis – via the mevalonate pathway using the Mevalonate-5P to Isopentenyl-PP route (Figure 8d). The Mevalonate-5P pathway is used in most higher eukaryotes as compared to the Mevalonate-3P pathway in Archaea and the MEP/DOXP pathway in bacteria, some plants and apicomplexan protists (Breitmaier 2006; Degenhardt et al. 2009; Eisenreich et al. 2004; Miziorko 2011; Oldfield and Lin 2012; Trapp and Croteau 2001). This likely excludes any horizontal gene transfer of mevalonate pathway genes as *Archegozetes* uses enzymes similar to those of other animals.

The biosynthesis of monoterpenes not only depends on very widespread enzymes, but also requires more specific enzymes downstream of GPP (Breitmaier 2006; Degenhardt et al. 2009; Trapp and Croteau 2001). For instance, Carpoglyphus lactis expresses a unique geraniol dehydrogenase (GeDH) – catalyzing the oxidation of geraniol to geranial – different from all previously characterized geraniol-related and alcohol dehydrogenases (ADHs) of animals and plants (Noge et al. 2008). We used the functionally validated Carpoglyphus-GeDH (Noge et al. 2008), blasted its sequence against the *Archegozetes* OGS and found a homologous sequence. We used both mite sequences in an alignment with plant, fungal and bacterial GeDHs and animal ADHs and constructed a maximum likelihood phylogeny (Figure 8e). Similar to the previous analysis including only Carpoglyphus-GeDH, we found that the Al-GeDH represent a new class of geraniol dehydrogenases different from those in plants, fungi or bacteria and not nested within animal ADHs (Figure 8e). This is why we hypothesize that Al-GeDH is a novel expansion of the geraniol dehydrogenases gene family and has not been acquired by horizontal gene transfer, like other biosynthesis and digestive enzymes in *Archegozetes* (Figure 7; (Brückner et al. 2020)).

Based on our mass spectrometry data of stable isotopes and genomic analysis, we propose that the following biochemical pathway leading to monoterpenes is of oribatid mites (Figure 8f and g): geraniol is likely to be synthesized from GPP – the universal precursor of all monoterpenes – either enzymatically by a geraniol synthase (GES) or a diphosphate phosphatase (DPP), but possibly also endogenously by dephosphorylation of GPP (Beran et al. 2019; Liu et al. 2015; Oswald et al. 2007; Zhou et al. 2014). For *Archegozetes*, we could not find any GES or specific DPP in the OGS, thus geraniol might be formed from GPP via endogenous dephosphorylation, but further research is required to verify or falsify this hypothesis. Subsequently, geraniol is oxidized to geranial by the pervious described GeDH (Figure 8e) and readily isomerized to neral. Trace amounts of geranial have been found in *Archegozetes* and it is common among other oribatid and astigmatid mites, supporting this idea (Koller et al. 2012; Kuwahara 2004; Raspotnig et al. 2008; Raspotnig et al. 2004). Also, there is no evidence that geraniol is converted into nerol, or that neral is formed directly via oxidation of nerol (Morita et al. 2004; Noge et al. 2005; Noge et al. 2008). The most parsimonious explanation for neryl formate synthesis would be an esterification of the corresponding terpene alcohol nerol. There is, however, no evidence of nerol in the traces of any oribatid or astigmatid mite species (Kuwahara 2004; Raspotnig 2009; Raspotnig et al. 2011). Aliphatic non-terpene formats in Astigmata are synthesized by dehomologation and generation of a one-carbon–shorter primary alcohol from an aldehyde via hydrolysis of formate in a biological Baeyer–Villiger oxidation catalyzed by a novel, uncharacterized enzyme (Shimizu et al. 2017). A similar reaction to synthesize terpene formates is unlikely, as the terpenoid backbone would be shortened by one-carbon and this does not happen in any possible scenario. The discovery of this Baeyer–Villiger oxidation mechanism, however, highlights the probability that there are many very unusual reactions that remain to be discovered in oribatid mites (Brückner et al. 2020).

## Conclusion

The integrated genomic and transcriptomic resources presented here for *Archegozetes longisetosus* allowed a number of insights into the molecular evolution and basic biology of decomposer soil mites. Our analysis of an oribatid mite genome also provides the foundation for experimental studies building on the long history of *Archegozetes*’ as a chelicerate model organism, which now enters the molecular genetics era (Aoki 1965; Heethoff et al. 2013; Norton et al. 1993; Palmer and Norton 1992). This includes the study of biochemical pathways, biochemistry, neuroethological bases of food searching behavior, and environmental impacts on genomes of complex, clonal organisms.

Our evolutionary comparisons across the Chelicerata revealed interesting patterns of genome evolution and how horizontal gene transfer might have shaped the feeding mode of soil mites. We also showed how oribatid glandular biology and chemical ecology are reflected in the genome. The community of researchers studying the fundamental biology of oribatid and other free-living, non-parasitic mites is growing. We think that providing these genomic and transcriptomic resources can foster a community effort to eventually transform basic research on these mites into a modern, molecular discipline.

Key priorities for a future community research effort include i) sequencing organ-specific transcriptomic data, ii) developing tools for genetic interrogation (RNAi or CRISPR/CAS9), iii) establishing reporter linages with germ-line stable modifications (e.g. GAL4/UAS misexpression systems), iv) constructing an whole-animal single-cell RNAseq expression atlas, and v) gathering more genomic data to improve the genome assembly. Please do not hesitate to contact the corresponding author, if you want to start your own culture of *Archegozetes*. He will be happy to provide you with starter specimens and share rearing protocols with you.

## Materials and Methods

### Mite husbandry

The lineage ‘ran’ (Heethoff et al. 2013) of the pantropical, parthenogenetic oribatid mite *Archegozetes longisetosus* was used in this study. Stock cultures were established in 2015 from an already existing line and fed with wheat grass (Triticum sp.) powder from Naturya. Cultures were maintained at 20-24°C and 90% relative humidity. Sterilized water and 3-5 mg wheat grass were provided three times each week.

### DNA extraction and Illumina sequencing

For the short-read library, DNA was extracted from ∼200 mites that were taken from the stock culture, starved for 24 h to avoid possible contamination from food in the gut, subsequently washed with 1% SDS for 10 sec. For extraction of living specimens, we used the Quick-DNA Miniprep Plus Kit (Zymo Research) according to the manufacturer’s protocol. Amounts and quality of DNA were accessed with Qubit dsDNA HS Kit (ThermoFisher) and with NanoDrop One (ThermoFisher) with target OD 260/280 and OD 260/230 ratios of 1.8 and 2.0-2.2, respectively. Extracted DNA was shipped to Omega Bioservices (Norcross, GA, USA) on dry ice for library preparation and sequencing. DNA library preparation followed the KAPA HyperPrep Kit (Roche) protocol (150 bp insert size) and 200 million reads were sequenced as 150bp paired-end on a HighSeq4000 (Illumina) platform.

### High-molecular weight DNA isolation and Nanopore sequencing

Genomic DNA was isolated from ∼300-500 mites starved for 24 h using QIAGEN Blood & Cell Culture DNA Mini Kit. Briefly, mites were flash frozen in liquid nitrogen and homogenized with a pestle in 1 ml of buffer G2 supplemented with RNase A and Proteinase K at final concentrations of 200 ng/µl and 1 µg/µl, respectively. Lysates were incubated at 50°C for 2 h, cleared by centrifugation at 5 krpm for 5 min at room temperature and applied to Genomic tip G/20 equilibrated with buffer QBT. Columns were washed with 4 ml of buffer QC and genomic DNA was eluted with 2 ml of buffer QF. DNA was precipitated with isopropanol, washed with 70% EtOH and resuspended in 50 µl of buffer EB. DNA was quantified with Qubit dsDNA HS Kit (ThermoFisher) and the absence of contaminants was confirmed with NanoDrop One (ThermoFisher) with target OD 260/280 and OD 260/230 ratios of 1.8 and 2.0-2.2, respectively. DNA integrity was assessed using Genomic DNA ScreenTape kit for TapeStation (Agilent Technologies).

Libraries for nanopore sequencing were prepared from 1 µg of genomic DNA using 1D Genomic DNA by Ligation Kit (Oxford Nanopore) following manufacturer’s instructions. Briefly, unfragmented DNA was repaired and dA tailed with a combination of NEBNext FFPE Repair Mix (New England Biolabs) and NEBNext End repair/dA-tailing Module (New England Biolabs). DNA fragments were purified with Agencourt AMPure XP beads (Beckman Coulter) and Oxford Nanopore sequencing adapters were ligated using NEBNext Quick T4 DNA Ligase (New England Biolabs). Following AMPure XP bead cleanup, ∼500 ng of the library was combined with 37.5 µL of SQB sequencing buffer and 25.5 µl of loading beads in the final volume of 75 µl and loaded on a MinION Spot-ON Flow Cell version R9.4 (Oxford Nanopore). Two flow cells were run on MinION device controlled by MinKNOW software version 3.1.13 for 48 hours each with local basecalling turned off generating 9.7 and 5.1 GB of sequence data. Post run basecalling was performed with Guppy Basecalling Software, version 3.4.5 (Oxford Nanopore). After filtering low quality reads (Q<7), the combined output of the two runs was 13.69 GB and 4.7 million reads.

### Genome assembly and contamination filtering

Read quality was assessed using FastQC v0.11.8 (Andrews 2010). Illumina adapters, low-quality nucleotide bases (phred score below 15) from the 3’ and 5’ ends and reads shorter than 50 bp were removed using cutadapt v1.18 (Martin 2011). From the filtered reads, in silico genome size estimates were calculated using k-mer based tools kmergenie v.1.7048 (Chikhi and Medvedev 2014), GenomeScope v1.0 (Vurture et al. 2017), and findGSE v0.1.0 R package (Sun et al. 2018). The latter two required a k-mer histogram computed by jellyfish v2.2.10 (Marçais and Kingsford 2011) with k-mer size of 21. The genome was assembled using 4.7 million long reads from two MinION runs (60x coverage) using Canu v1.8 with default settings and setting the expected genome size to 200 Mb (Koren et al. 2017). To improve assembly quality, paired end Illumina reads were mapped to the genome with BWA aligner (Li and Durbin 2009) using BWA-MEM algorithm and polished with Pilon v. 1.23 with ‘—changes’ and ‘--fix all’ options (Walker et al. 2014). Assembled contigs identified as bacterial and fungal contaminants based on divergent GC content from most *Archegozetes* contigs, high coverage and blast homology to the nt database (downloaded February 2019, Evalue 1e^-25^) were removed using Blobtools v1.0 (Laetsch and Blaxter 2017).

### Identification, classification and masking of repetitive element

Repetitive elements in the genome *Archegozetes* were identified using a species-specific library generated with RepeatModeler v 1.0.11 (Bao et al. 2015; Smit and Hubley 2008) and MITE tracker (Crescente et al. 2018) and annotated by RepeatClassifier, a utility of the RepeatModeler software that uses the RepBase database (version Dfam_Consensus-20181026). Unclassified repeat families from both programs were run through CENSOR v 4.2.29 (Kohany et al. 2006) executable censor.ncbi against the invertebrate library v 19.03 to provide further annotation. Predicted repeats were removed if they had significant blast homology (E-value 1e^-5^) to genuine proteins in the NCBI nr database and/or a local database of arthropod genomes (*Drosophila melanogaster, Tribolium castaneum, Tetranychus urticae, Leptotrombidium deliense, Dinothrombium tinctorium, Sarcoptes scabiei, Euroglyphus maynei, Galendromus occidentalis, Dermatophagoides pteronyssinus*). Unclassified repeats with blast homology to known TEs were retained whereas those with no blast homology were removed (Petersen et al. 2019). The remaining repeat families were combined with the Arthropoda sequences in RepBase and clustered using vsearch v 2.7.1 (--iddef 1 --id 0.8 --strand both; (Rognes et al. 2016)). The filtered repeat library was used to soft mask the *A. longisetosus* using RepeatMasker v 4.07 (Smit et al. 1996-2010).A summary of the masked repeat content was generated using the “buildSummary.pl” script, the Kimura sequence divergence calculated using the “calcDivergenceFromAlign.pl” script and the repeat landscape visualized using the “createRepeatLandscape.pl” script, all utilities of RepeatMasker.

### Gene prediction and annotation

Both ab inito and reference-based tools were used for gene prediction using modified steps of the funannotate pipeline (Palmer and Stajich 2017). The *ab inito* tool GeneMark-ES v4.33 (Ter-Hovhannisyan et al. 2008) was used along with reference based tools BRAKER v2.1.2 (Bruna et al. 2020) using RNAseq reads discussed below and PASA v 2.3.3 (Haas et al. 2008) using genome-guided transcriptome assembly from Trinity described below. Lastly, *Tetranychus urticae* gene models from the NCBI database (GCF_000239435.1) were aligned to the contigs using GeMoMa (Keilwagen et al. 2019). All gene predictions were combined in EvidenceModeler (Haas et al. 2008) with the following weights: GeMoMa =1, PASA = 10, other BRAKER = 1, and GeneMark = 1. Predicted tRNAs using tRNAscan-SE v 2.0.3 (Chan and Lowe 2019) were combined with the gene predictions in the final gene feature format (GFF) file and filtered for overlap using bedtools (Quinlan and Hall 2010) intersect tool (Quinlan and Hall 2010).

The predicted genes were searched against the NCBI nr (February 2019) (Pruitt et al. 2005), SwissProt (February 2019) (Bairoch and Apweiler 2000), a custom-made Chelicerata database including genomes of *Tetranychus urticae, Leptotrombidium deliense, Dinothrombium tinctorium, Sarcoptes scabiei, Euroglyphus maynei, Galendromus occidentalis, Metaseiulus occidentalis, Dermatophagoides pteronyssinus, Trichonephila clavipes, Stegodyphus mimosarum, Centruroides sculpturatus, Ixodes scapularis and Parasteatoda tepidariorum* (all downloaded Feb 2019), PFAM (v 32, August 2018) (Bateman et al. 2004), merops (v 12, October 2017) (Rawlings et al. 2010) and CAZY (v 7, August 2018) (Cantarel et al. 2009) databases. The results of the hmm-based (Eddy 2011) PFAM and CAZY searches were filtered using cath-tools v 0.16.2 (https://cath-tools.readthedocs.io/en/; E-value 1e^-5^) and the blast-based searches were filtered by the top hit (E-value 1e^-5^ threshold). Predicted genes were also assigned to orthologous groups using eggNOG-mapper (Huerta-Cepas et al. 2017). Gene annotation was prioritized by the SwissProt hit if the E-value < 1e^-10^ followed by NCBI annotation, the custom Chelicerata database and if no homology was recovered, then the gene was annotated as, “hypothetical protein”. Final annotation was added to the GFF file using GAG (Geib et al. 2018).

### Analysis of the official gene set (OGS)

To allow the OGS to be used as resources for functional studies, we assigned functional categories based on Gene Ontology (GO) and the Kyoto Encyclopedia of Genes and Genomes (KEGG) (Consortium 2004; Kanehisa and Goto 2000). GO terms for the respective genes models of the OGS were assigned based on the gene id with highest homology from the SwissProt database or NCBI nr database (Bairoch and Apweiler 2000; Pruitt et al. 2005). A custom database of GO terms was created with makeOrgPackage function in the R package AnnotationForge v1.26.0 (Carlson and Pagès 2019). Over-representation analysis of GO terms was tested using the enrichGO function in the R package clusterProfiler v3.12.0 (Yu et al. 2012) with a hypergeometric distribution and a Fisher’s Exact test. P-values were adjusted for multiple comparisons using false discovery rate correction (Benjamini and Hochberg 1995). Resulting enriched GO terms were processed with GO slim (Consortium 2019) and the final list of over represented GO terms was used to plot the number of genes in a respective category.

KEGG orthology terms were assigned from single-directional best hit BLAST searches of each gene model on the KEGG Automatic Annotation Server (Moriya et al. 2007). Additionally, we ran GhostKOALA (Kanehisa et al. 2016) (GHOSTX searches for KEGG Orthology And Links Annotation) to obtain KEGG orthology terms. Compared to conventional BLAST searches, GhostKOALA is about 100 times more efficient than BLAST to remote homologs by using suffix arrays (Suzuki et al. 2014).

### Orthology and phylogenomic analyses

Orthologs of *A. longisetosus*, other species within Acari, Chelicerata and the fruit fly *Drosophila* were identified using OrthoFinder v 2.3.3 (-M msa –A mafft –T fasttree; (Emms and Kelly 2015a)). Prior to running OrthoFinder, isoform variants were removed from the gene predictions using CD-Hit (Fu et al. 2012). Trees of orthogroups with at least 80% of taxa present (n= 4,553) were constructed using fasttree v 2.1.10 (Price et al. 2010), trimmed with TrimAl v 1.4.1 (-keepheader -fasta –gappyout; (Capella-Gutiérrez et al. 2009)) and paralogs pruned using phylotreepruner v 1.0 (min_number_of_taxa =18, bootstrap_cutoff= 0.7, longest sequence for a given orthogroup=u; (Kocot et al. 2013)). Alignments shorter than 100 amino acids were removed, leaving 1,121 orthogroups.

For the maximum likelihood analysis, the trimmed and pruned alignments were concatenated into a supermatrix using FasConCat v1.04 (Kück and Meusemann 2010) composed of 377,532 amino acids and the best substitution models determined using PartitionFinder v 2.1.1 (Lanfear et al. 2016). The maximum likelihood consensus phylogeny from the supermatrix and partition scheme was constructed using IQ-tree and 1,000 ultrafast bootstrap replicates (Nguyen et al. 2015). For the coalescence species tree reconstruction, gene trees were generated using IQ-tree v 1.6.12 on the trimmed alignments of the 1,121 filtered orthogroups and processed using ASTRAL v 5.6.3 (Zhang et al. 2018). Branch lengths are presented in coalescent units (differences in the 1,121 gene trees) and the node values reflect the local posterior probabilities.

### RNA sequencing and transcriptome assembly

For RNA extraction, about 200 mites of all life stages were taken from stock culture and subsequently washed with 1% SDS for 10 s. RNA was extracted from living specimens using the Quick-RNA MiniPrep Kit (Zymo Research) according to the manufacturer’s protocol. Quantity and quality of RNA were accessed using a Qubit fluorometer and NanoDrop One (Thermo Fisher Scientific), respectively.

Extracted RNA was shipped to Omega Bioservices (Norcross, GA, USA) on dry ice for library preparation and sequencing. Whole animal RNA was used for poly-A selection, cDNA synthesis and library preparation following the Illumina TruSeq mRNA Stranded Kit protocol. The library was sequenced with 100 million 150 bp paired-end on a HighSeq4000 platform. For the genome-guided assembly of the transcriptome a bam-file was created from the genome using STAR (Dobin et al. 2013). RNAseq reads were in silico normalized and subsequently used together with the bam-file to assemble the transcripts using Trinity v2.8.4 (Grabherr et al. 2011; Haas et al. 2013), yielding an assembly with a total length of 162.8 Mbp, an N50= 2994 bp and a BUSCO score (Simão et al. 2015) of C:96.3% [S:36.5%,D:59.8%], F:1.3%, M:2.4%.

### Life-stage specific RNAseq

For life-stage specific RNAseq, we collected 15 specimens per life stage from the stock culture that were split into three replicates of five individuals. Whole animals (for all stages but eggs) were flash frozen in 50 µl TRIzol using a mixture of dry ice and ethanol (100%) and stored at −80°. RNA was extracted using a combination of the TRIzol RNA isolation protocol (Life Technologies) and RNeasy Mini Kit (Qiagen) (Kitchen et al. 2015). The TRIzol protocol was used for initial steps up to and including the chloroform extraction. Following tissue homogenization, an additional centrifugation step was performed at 12,000 × g for 10 min to remove tissue debris. After the chloroform extraction, the aqueous layer was combined with an equal volume of ethanol and the RNeasy Mini Kit was used to perform washes following the manufacturer’s protocol. Eggs were crushed using pipette tips and directly stored in a mixture of cell lysis buffer and murine RNase Inhibitor (New England Biolab).

We used the NEBNext® Single Cell/Low Input RNA Library Prep Kit for Illumina® together with NEBNext® Multiplex Oligos for Illumina® (New England Biolab) for library preparation, including reverse transcription of poly(A) RNA, amplification full-length cDNA, fragmentation, ligation and final library amplification according to the manufacturer’s protocol. We performed cDNA amplification for 16 (18 for egg samples) PCR cycles and final library amplification 8 PCR cycles. In total, we constructed 18 libraries (three for each life stage). The quality and concentration of the resulting libraries were assessed using the Qubit High Sensitivity dsDNA kit (Thermo Scientific) and Agilent Bioanalyzer High Sensitivity DNA assay. Libraries were sequenced on an Illumina HiSeq2500 platform (single-end with read lengths of 50 bp) with ∼18 million reads per library.

Illumina sequencing reads were pseudoaligned to the bulk transcriptome and quantified (100 bootstrap samples) with kallisto 0.46.0 (Bray et al. 2016) using default options for single-end reads. Fragment length sizes were extracted from the Agilent Bioanalyzer runs. For life-stage specific differential expression analysis, kallisto quantified RNAseq data was processes with sleuth 0.30.0 (Pimentel et al. 2017) using Likelihood Ratio tests in R 3.6.1 (R_Core_Team 2019). The average transcripts per million (tpm) values for each target transcript were extracted from the sleuth object (see R script) and used with the Heatmapper tool (Babicki et al. 2016) to produce an unclustered heatmap showing relative expression levels. UpSetR (Conway et al. 2017) was used to compare the number of unique and shared expressed genes across life stages.

### Horizontal gene transfer events identification

To detect HGTs, we used the published tool “./Lateral_gene_transfer_predictor.py” (Thorpe et al. 2018) to calculate the Alien Index described by (Gladyshev et al. 2008) and (Flot et al. 2013). All predicted genes were compared to the NCBI nr database as previously described (Thorpe et al. 2018). Results to Arthropoda (tax id 6656) were ignored in the downstream calculations. The HGT candidates were filtered for contamination identified by both Blobtools (Laetsch and Blaxter 2017) and the Alien Index (AI > 30 and >70% percent identity to a non-metazon sequence). The candidates were further filtered for > 50% overlap with predicted repeats using the bedtools intersect tool with the RepeatMasker gff file and expression from any developmental stage. Introns were scored manually from visualization in IGV genome browser (Robinson et al. 2011) and GC content for all predicted genes was calculated using the bedtools nuc tool.

### Analysis of chemosensory and photoreceptor gene families

The search and analysis chemosensory genes largely followed the procedure outlined by Dong et al. (Dong et al. 2018) with slight modifications. First, the *Archegozetes* official gene set (OGS) was searched using BLASTP (E-value, <1 × 10^−3^) against the following queries for the different chemosensory gene families. The OGS was queried against i) *D. melanogaster, D. mojavensis, Anopheles gambiae, Bombyx mori, T. castaneum, Apis mellifera, Pediculus humanus humanu*s, and *Acyrthosiphon pisum* odorant binding proteins (OBPs) (Vieira and Rozas 2011); ii) *D. melanogaster*, *D. mojavensis*, *A. gambiae*, *B. mori*, *T. castaneum*, *A. mellifera*, *P. humanus humanus*, *A. pisum*, *I. scapularis*, and *Daphnia pulex* small chemosensory proteins (CSP) (Niimura and Nei 2005; Robertson and Wanner 2006; Vieira and Rozas 2011); iii) *D. melanogaster* and *A. mellifera* odorant receptors (Niimura and Nei 2005; Robertson and Wanner 2006); iv) *D. melanogaster, A. mellifera, I. scapularis, T. urticae, T. mercedesae*, and *M. occidentalis* gustatory receptors (GRs) (Dong et al. 2017; Gulia-Nuss et al. 2016; Hoy et al. 2016; Ngoc et al. 2016; Robertson and Wanner 2006; Robertson et al. 2003); v) a comprehensive list of iGluRs and IRs across vertebrates and invertebrates (Croset et al. 2010), as well as those identified in the *T. mercedesae, D. tinctorium* and *L. deliense* genome projects (Dong et al. 2017; Dong et al. 2018). Second, all candidate *Archegozetes* sequences were reciprocally blasted (BLASTP, E-value <1 × 10^−3^) against the NCBI database (Pruitt et al. 2005) and all sequences that did not hit one of the respective receptors or transmembrane proteins were removed from the list. Third, for phylogenetic analysis of IRs and GRs from *Archegozetes* were aligned with IRs from *D. melanogaster, T. urticae, D. tinctorium* and *L. deliense* and GRs from iv) *D. melanogaster, T. mercedesae, I. scapularis*, and *M. occidentalis*, respectively, using MAFFT (v 7.012b) with default settings (Katoh and Standley 2013). Poorly aligned and variable terminal regions, as well as several internal regions of highly variable sequences were excluded from the phylogenetic analysis. Fourth, maximum likelihood trees were constructed with the IQ-TREE pipeline (v 1.6.12) with automated model selection using 1,000 ultrafast bootstrap runs (Nguyen et al. 2015).

Reference opsin genes and opsin-like sequences were obtained from Dong et al. (Dong et al. 2018) and used to query the *Archegozetes* OGS using BLASTP (E-value, <1 × 10^−5^). Subsequently, candidates sequenced were reciprocally blasted against NCBI using the same settings and only retained if they hit an opsin or opsin-like gene. The *Archegozetes* candidates were aligned with the query sequence list using MAFFT (v 7.012b) with default settings (Katoh and Standley 2013). This opsin gene alignment phylogenetically analyzed using the IQ-TREE pipeline (v 1.6.12) with automated model selection and 1,000 ultrafast bootstrap runs (Nguyen et al. 2015).

### Gene family phylogenies

We used the following workflow to analyses genes related to Figure 5 (hox and developmental genes), Figure 7 (cell wall-degrading enzyme encoding genes) and Figure 8 (alcohol and geraniol dehydrogenases genes). Generally, protein orthologs were retrieved from NCBI (Pruitt et al. 2005), and aligned using MUSCLE (Edgar 2004) or MAFFT (v 7.012b) (Katoh and Standley 2013) and ends were manually inspected and trimmed. The resulting final protein sequence alignments used to construct a maximum likelihood (ML) phylogenetic tree with either i) PhyML with Smart Model Selection (Guindon et al. 2010; Lefort et al. 2017) or ii) the IQ-TREE pipeline with automated model selection (Nguyen et al. 2015). The ML trees were constructed using either 1,000 ultrafast bootstrap runs (IQ-TREE) or approximate-likelihood ratio test (PhyML) was used to assess node support.

### Feeding experiments with labelled precursors and chemical analysis (GC/MS)

Stable isotope incorporation experiments were carried out as previously described (Brückner et al. 2020). Briefly, mites were fed with wheat grass containing a 10% (w/w) mixture of three antibiotics (amoxicillin, streptomycin and tetracycline) and additionally, we added 25% (w/w) of the stable isotope-labelled precursors [^13^C_6_] D-glucose (Cambridge Isotope Laboratories, Inc.) as well as a control with untreated wheat grass. Cultures were maintained for one generation and glands of adult specimens were extracted one week after eclosion by submersing groups of 15 individuals in 50 µl hexane for 5 min, which is a well-established method to obtain oil gland compounds from mites (Brückner and Heethoff 2016; 2017; Brückner et al. 2017b; Raspotnig et al. 2008).

Crude hexane extracts (2-5 µl) were analysed with a GCMS-QP2020 gas chromatography – mass spectrometry (GCMS) system from Shimadzu equipped with a ZB-5MS capillary column (0.25 mm x 30m, 0.25 µm film thickness) from Phenomenex. Helium was used a carrier gas with a flow rate of 2.14 ml/min, with splitless injection and a temperature ramp was set to increase from 50°C (5 min) to 210°C at a rate of 6°C/min, followed by 35°C/min up to 320°C (for 5 min). Electron ionization mass spectra were recorded at 70 eV and characteristic fragment ions were monitored in single ion mode.

## Acknowledgment

We thank Joe Parker for making his laboratory space and resources available to us. Michael Heethoff, Sebastian Schemlzle, Benjamin Weiss and Martin Kaltenpoth graciously allowed us to use some of their unpublished images. Roy A. Norton provided invaluable comments to the manuscript and collected the first specimens of *Archegozetes longisetosus* giving rise to the current laboratory strain. This work was supported by a grant from the Caltech Center for Environmental Microbial Interactions (CEMI) to AB. AB is Simons Fellow of the Life Sciences Research Foundation (LSRF).

## Ethics statement

There are no legal restrictions on working with mites.

## Authors contributions

AB had the initial idea for the study; AB, AAB and SAK design research; IAA performed long-read sequencing and assembled the genome; AB performed all other experimental work; AAB analyzed hox and life-stage specific expression data; AB analyzed chemical data; SAK and AB performed bioinformatic analyses; AB wrote the first draft of the manuscript with input from AAB and SAK; SAK revised the manuscript. All authors gave final approval for publication.

## Data availability

Genomic and transcriptomic data generated for his project can be found on NCBI under the accession numbers PRJNA683935 and PRJNA683999. All other data related to this manuscript can be found at https://doi.org/10.22002/D1.1876 under a cc-by-nc-4.0 license (Brückner 2021).

## Supplementary Material

### Supplementary Figures

**Figure S1** Results of *in silico* genome size estimations based on jellyfish k-mer counting using a: GenomeScope v1.0 and b and c: the findGSE v0.1.0 R package (Sun et al. 2018).

**Figure S2** Phylogenetic placement of *Archegozetes longisetosus* among other chelicerates. a: Maximum likelihood phylogeny based on concatenation of 1,121 orthologs Branch lengths unit is substitutions per site and the node values reflect bootstrap supports. b: Coalescence species tree reconstruction of the 1,121 filtered orthogroups. Branch lengths are presented in coalescent units (differences in the 1,121 gene trees) and the node values reflect the local posterior probabilities.

**Figure S3** Comparisons of protein-coding genes of 23 arthropod species, including *Archegozetes*. The bar charts show the proportion of orthrogroup conservation with each species (see insert legend) based on OrthoFinder clustering.

**Figure S4** Maximum likelihood phylogenetic analyses of the *A. longisetosus* Paired protein orthologs. (a) Maximum likelihood tree showing the relationship of the Eyegone, Pax-3/7, and Pax-6 clades as collapsed subtrees. (b) The un-collapsed clade in A showing the phylogenetic relationships of selected Eyegone proteins and the putative *A. longisetosus* Eyegone ortholog. (c) The un-collapsed clade in A showing the phylogenetic relationships of selected Pax-3/7 proteins and the putative *A. longisetosus* Pax-3/7 orthologs. (d) The un-collapsed clade in A showing the phylogenetic relationships of selected Pax-6 proteins and the putative *A. longisetosus* Pax-6 ortholog. All *A. longisetosus* orthologs are in red, and the *D. melanogaster* orthologs are in blue. Node support was calculated using the approximate likelihood ratio (aLRT) method and is represented by the color of each node. All taxa are represented by their species names, gene names if given, and their NCBI accession numbers.

**Figure S5** Maximum likelihood phylogenetic analyses of the *A. longisetosus* Eyes absent (Eya) protein ortholog and selected metazoan Eya proteins. The *A. longisetosus* ortholog is in red, and the *D. melanogaster* orthologs are in blue. Node support was calculated using the approximate likelihood ratio (aLRT) method and is represented by the color of each node. All taxa are represented by their species names, gene names if given, and their NCBI accession numbers.

**Figure S6** Maximum likelihood phylogenetic analyses of the *A. longisetosus* Hairy protein ortholog and selected metazoan Hairy proteins. Hairy/E(spl) proteins were used as an outgroup. The *A. longisetosus* ortholog is in red, and the *D. melanogaster* orthologs are in blue. Node support was calculated using the approximate likelihood ratio (aLRT) method and is represented by the color of each node. All taxa are represented by their species names, gene names if given, and their NCBI accession numbers.

**Figure S7** Maximum likelihood phylogenetic analyses of the *A. longisetosus* Omb, T-box H15, and TBX1 protein orthologs and selected metazoan T-box proteins. All *A. longisetosus* orthologs are in red, and the *D. melanogaster* orthologs are in blue. Node support was calculated using the approximate likelihood ratio (aLRT) method and is represented by the color of each node. All taxa are represented by their species names, gene names if given, and their NCBI accession numbers.

**Figure S8** Maximum likelihood phylogenetic analyses of the *A. longisetosus* Runt protein ortholog and selected arthropod Runt proteins. The *A. longisetosus* ortholog is in red, and the *D. melanogaster* orthologs are in blue. Node support was calculated using the approximate likelihood ratio (aLRT) method and is represented by the color of each node. All taxa are represented by their species names, gene names if given, and their NCBI accession numbers. See

**Figure S9** Maximum likelihood phylogenetic analyses of the *A. longisetosus* Six family protein orthologs and selected metazoan Six family proteins. All *A. longisetosus* orthologs are in red, and the *D. melanogaster* orthologs are in blue. Node support was calculated using the approximate likelihood ratio (aLRT) method and is represented by the color of each node. All taxa are represented by their species names, gene names if given, and their NCBI accession numbers.

**Figure S10** Maximum likelihood phylogenetic analyses of the *A. longisetosus* Sp-family protein orthologs and selected metazoan Sp-family proteins. All *A. longisetosus* orthologs are in red, and the *D. melanogaster* orthologs are in blue. Node support was calculated using the approximate likelihood ratio (aLRT) method and is represented by the color of each node. All taxa are represented by their species names, gene names if given, and their NCBI accession numbers.

**Figure S11** Maximum likelihood phylogenetic analyses of the *A. longisetosus* Wnt-family protein orthologs and selected metazoan Wnt proteins. The tree is organized as a cladogram for easier viewing. All *A. longisetosus* orthologs are in red, and the *D. melanogaster* orthologs are in blue. Node support was calculated using the approximate likelihood ratio (aLRT) method and is represented by the color of each node. All taxa are represented by their species names, gene names if given, and their NCBI accession numbers.

**Figure S12** Unrooted maximum-likelihood phylogenetic trees of cell-wall degrading enzymes based on the alignment of amino acid sequences. Branch lengths unit is substitutions per site and the node values reflect bootstrap supports. *Archegozetes* sequences are highlighted in red.

**Figure S13** Maximum likelihood phylogeny of ionotropic receptors and ionotropic glutamate receptors of *Archegozetes* (Along), *Dinothrombium* (Dt), *Leptothrombidium* (Ld), *Tetranychus* (Tu) and *Drosophila* (Dmel). The tree was rooted to the middle point. Branch lengths unit is substitutions per site and the node values reflect bootstrap supports.

**Figure S14** Maximum likelihood phylogenetic tree of gustatory receptors of *Archegozetes* (Along), *Ixodes* (Is), *Tropilaelaps* (Tm), *Metaseiulus* (Mocc) and *Drosophila* (Dmel). The tree was rooted to the middle point. Branch lengths unit is substitutions per site and the node values reflect bootstrap supports.

**Figure S15** Unrooted maximum-likelihood tree of glycoside hydrolase family 48 members (GH_48) from Streptomyces bacteria and HGT genes from other arthropods as well as *Archegozetes* (GH_48 II). Branch lengths unit is substitutions per site and the node values reflect bootstrap supports.

### Supplementary Table

Table S1 Contamination contigs identified from Blobtools.

Table S2 Phylogenetic statistics of the PhyML constructed trees as well as the matrices selected by the SMS model selection tool.

Table S3 The average transcript per million (tpm) values for the transcripts highlighted in the heatmap (Figure 5f) for each instar stage.

Table S4 Candidate HGTs identified from the *Archegozetes* genome. The genes were filtered first if they were predicted to be contamination from Blobtools and the Alien Index report, second if they overlapped predicted repeats by ≥ 50%, and third if they were not expressed in any developmental stage. Annotation is provided from similarity searches against the NCBI nr database, other oribatid mite and eggNOG database. The taxonomy of the sequences upstream and downstream of each candidate HGT was determine using the eggNOG predicted taxonomic group.

