## Supplemental Table S2 for "Molecular evolutionary trends and biosynthesis pathways in the Oribatida revealed by the genome of *Archegozetes longisetosus*" (from another version of this preprint: version 1, 2020-12-11)

| Tree | Log-likelihood | SMS Best Model |
| --- | --- | --- |
| Paired | -166943.2697 | VT +G+I+F |
| Eya | -60688.88985 | VT +G+F |
| Hairy | -63306.72113 | VT +G+I+F |
| Omb | -94052.6894 | VT +G+I+F |
| Runt | -27676.65469 | VT +G+I+F |
| Six Family | -50607.26768 | VT +G+I+F |
| Sp6-9 | -85523.56913 | VT +G+I+F |
| Wnt Family | -101311.944 | LG +G |
