## Supplemental Table S3 for "Molecular evolutionary trends and biosynthesis pathways in the Oribatida revealed by the genome of *Archegozetes longisetosus*" (from another version of this preprint: version 1, 2020-12-11)

| gene name | transcript ID | egg | larva | protonymph | deuteronymph | tritonymph | adult |
| --- | --- | --- | --- | --- | --- | --- | --- |
| *Distal-less* | TRINITY_GG_4702_c893_g1_i8 | **109.5** | 11.1 | 2.0 | 3.2 | 4.5 | 2.2 |
| *homothorax* | TRINITY_GG_4744_c51_g1_i1 | **4.0** | 9.5 | 1.0 | 2.2 | 2.4 | 11.1 |
| *extradenticle* | TRINITY_GG_4863_c311_g1_i1 | **15.3** | 3.7 | 1.5 | 1.0 | 2.2 | 2.1 |
| *dachshund* | TRINITY_GG_5120_c350_g1_i2 | **99.3** | 26.4 | 4.5 | 12.0 | 14.1 | 4.2 |
| *sp6-9* | TRINITY_GG_4863_c412_g1_i6 | **39.4** | 4.9 | 1.0 | 1.1 | 4.7 | 1.7 |
| *omb* | TRINITY_GG_4863_c1162_g1_i3 | **2.3** | 4.1 | 0.1 | 1.0 | 1.7 | 0.4 |
| *Pax6-A* | TRINITY_GG_2648_c164_g1_i2 | **1.0** | 7.9 | 1.1 | 2.1 | 0.7 | 0.3 |
| *Pax6-B* | TRINITY_GG_2648_c164_g1_i1 | **1.5** | 0.5 | 0.5 | 0.2 | 0.6 | 1.9 |
| *sine oculis* | TRINITY_GG_5990_c254_g1_i5 | **9.1** | 5.4 | 0.8 | 3.4 | 2.0 | 6.1 |
| *eya* | TRINITY_GG_4744_c173_g1_i10 | **0.7** | 0.4 | 0.1 | 0.0 | 0.1 | 1.2 |
| *eyegone* | TRINITY_GG_64_c64_g1_i12 | **0.7** | 0.6 | 0.5 | 0.5 | 0.5 | 1.4 |
| *six3/optix* | TRINITY_GG_5990_c859_g1_i1 | **34.3** | 5.9 | 1.5 | 1.8 | 3.4 | 2.5 |
| *orthodenticle* | TRINITY_GG_5990_c97_g1_i1 | **43.1** | 11.6 | 1.8 | 4.3 | 16.2 | 39.0 |
| *atonal* | TRINITY_GG_4863_c1996_g1_i1 | **15.0** | 6.8 | 6.2 | 5.4 | 4.3 | 4.4 |
| *engrailed* | TRINITY_GG_4702_c1211_g1_i1 | **32.6** | 302.7 | 16.7 | 18.6 | 7.4 | 3.8 |
| *hedgehog* | TRINITY_GG_4702_c448_g1_i24 | **1.0** | 4.3 | 1.3 | 1.3 | 3.2 | 0.5 |
| *wingless* | TRINITY_GG_4863_c585_g1_i7 | **17.6** | 6.9 | 9.0 | 5.6 | 4.2 | 0.0 |
| *patched* | TRINITY_GG_4702_c274_g1_i1 | **18.4** | 7.1 | 2.2 | 1.9 | 2.1 | 2.3 |
| *pax3/7* | TRINITY_GG_3318_c2184_g1_i1 | **0.4** | 0.9 | 0.1 | 0.0 | 0.0 | 2.4 |
| *runt* | TRINITY_GG_6648_c85_g1_i1 | **0.2** | 1.0 | 0.3 | 0.1 | 0.2 | 4.2 |
| *hairy* | TRINITY_GG_6485_c290_g1_i1 | **50.4** | 3.4 | 1.2 | 0.4 | 1.9 | 3.7 |
