## Supplementary figures and images for "Molecular evolutionary trends and biosynthesis pathways in the Oribatida revealed by the genome of *Archegozetes longisetosus*" (from another version of this preprint: version 1, 2020-12-11)

### Supplemental Figure S1

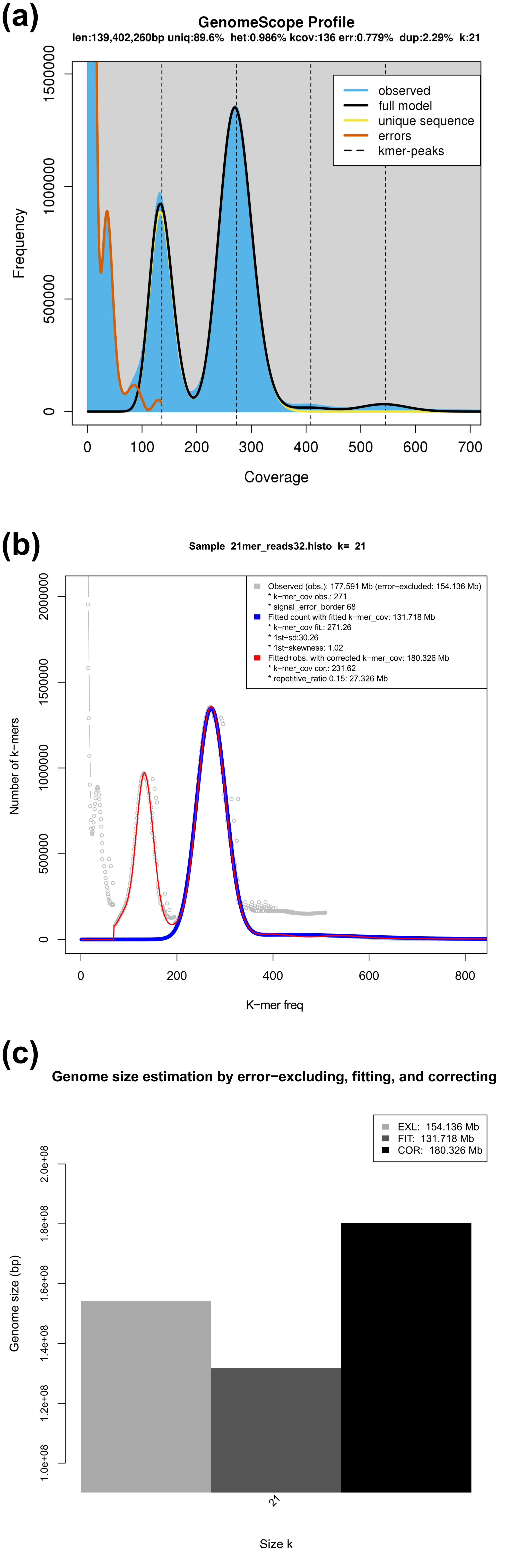

### Supplemental Figure S2

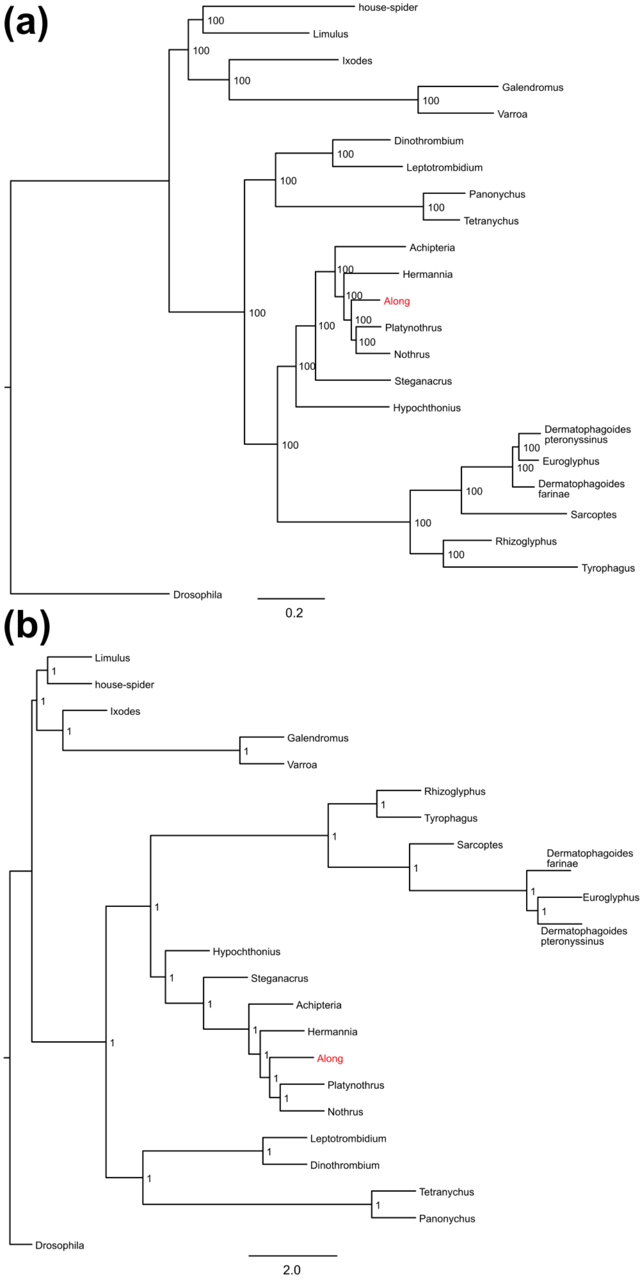

### Supplemental Figure S3

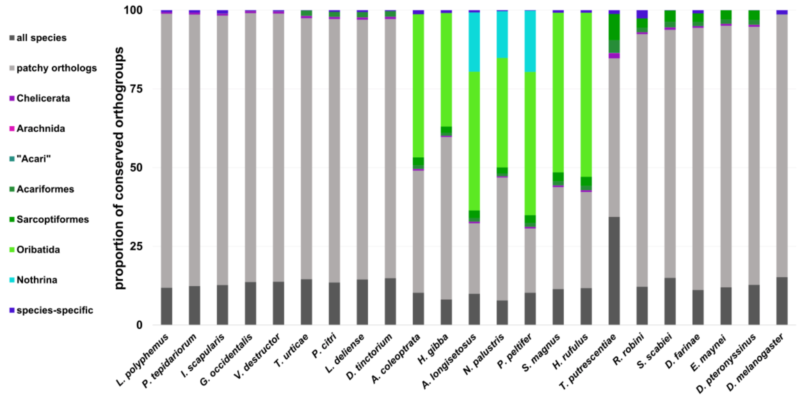

### Supplemental Figure S4A

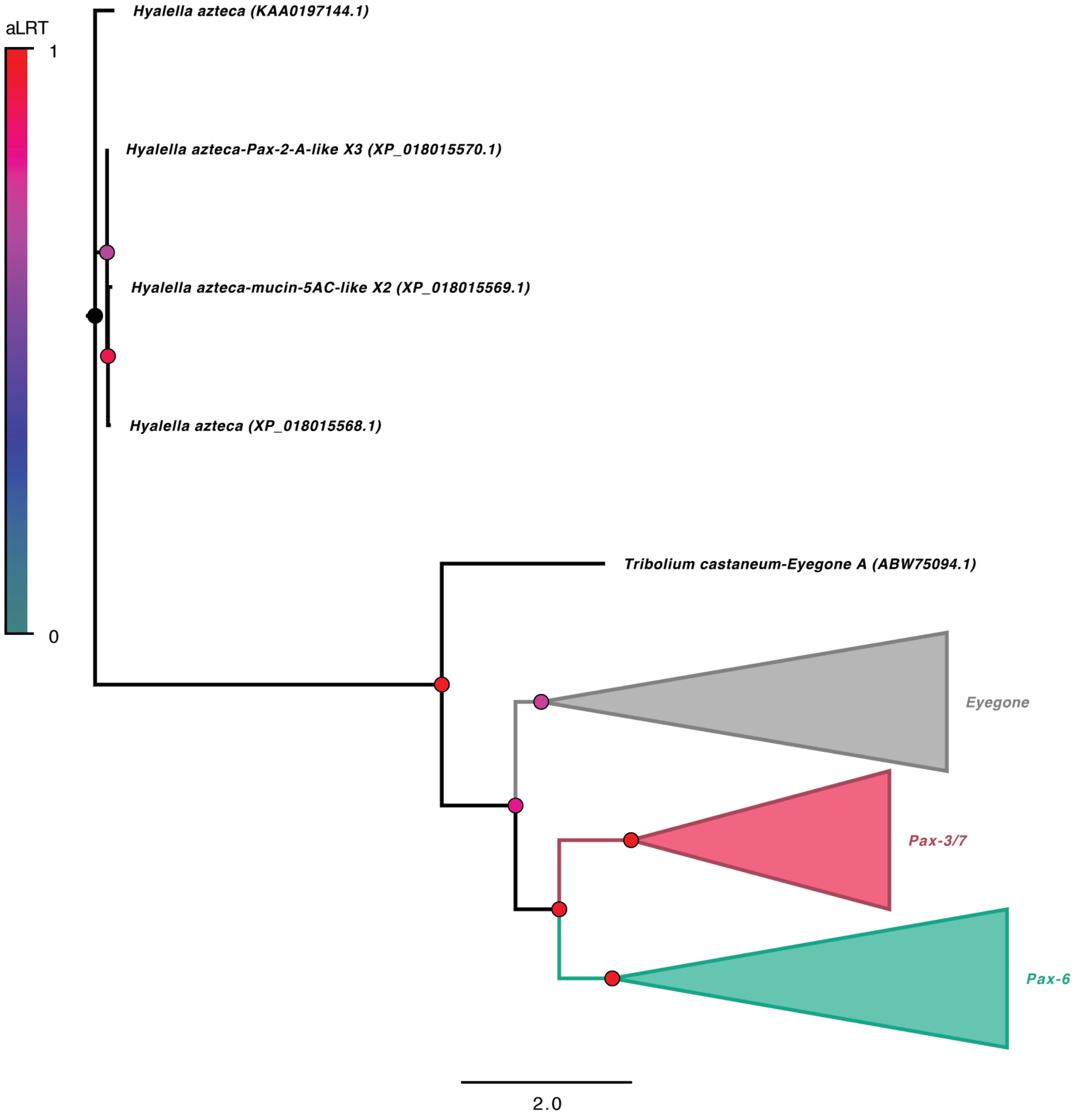

### Supplemental Figure S4B

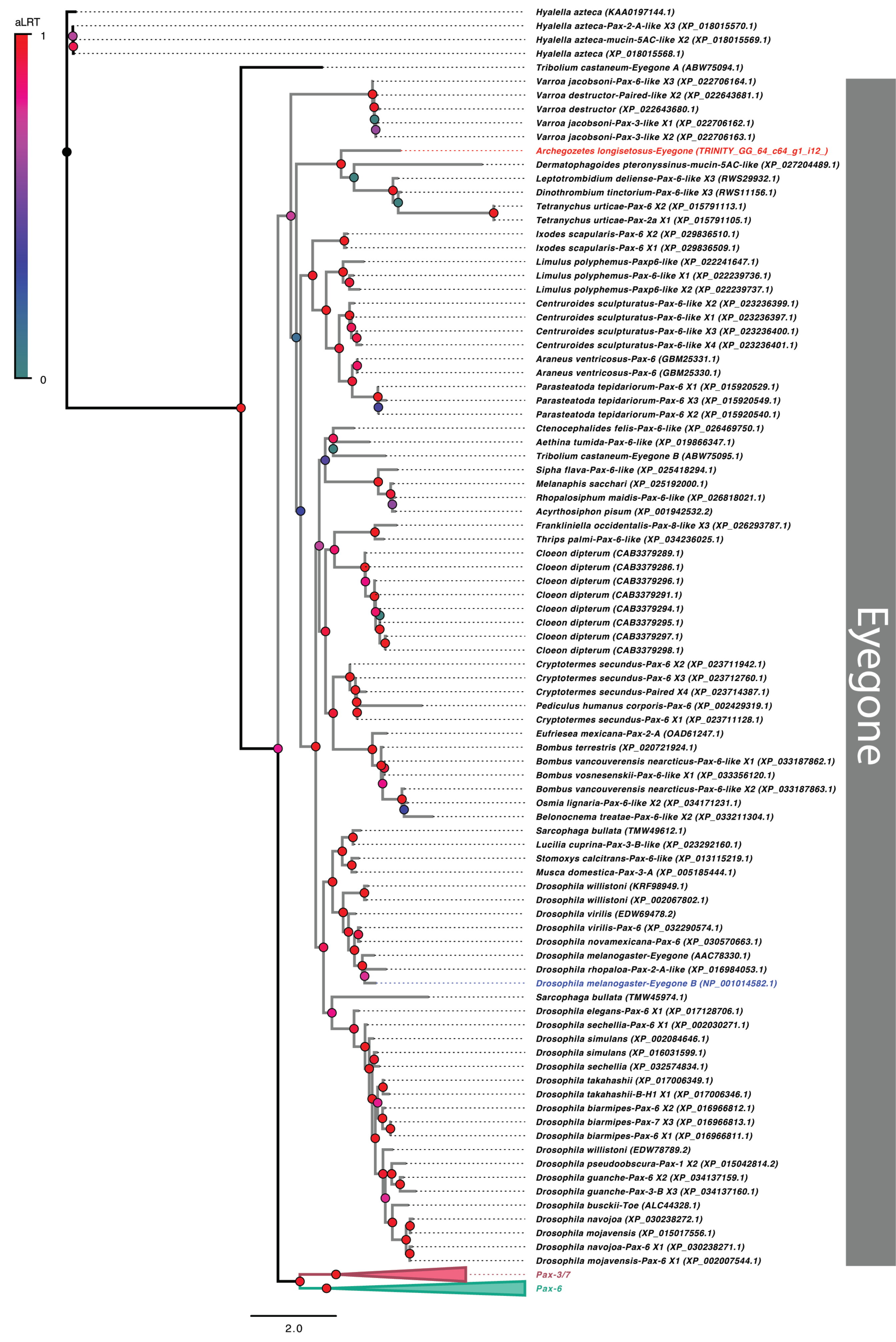

### Supplemental Figure S4C

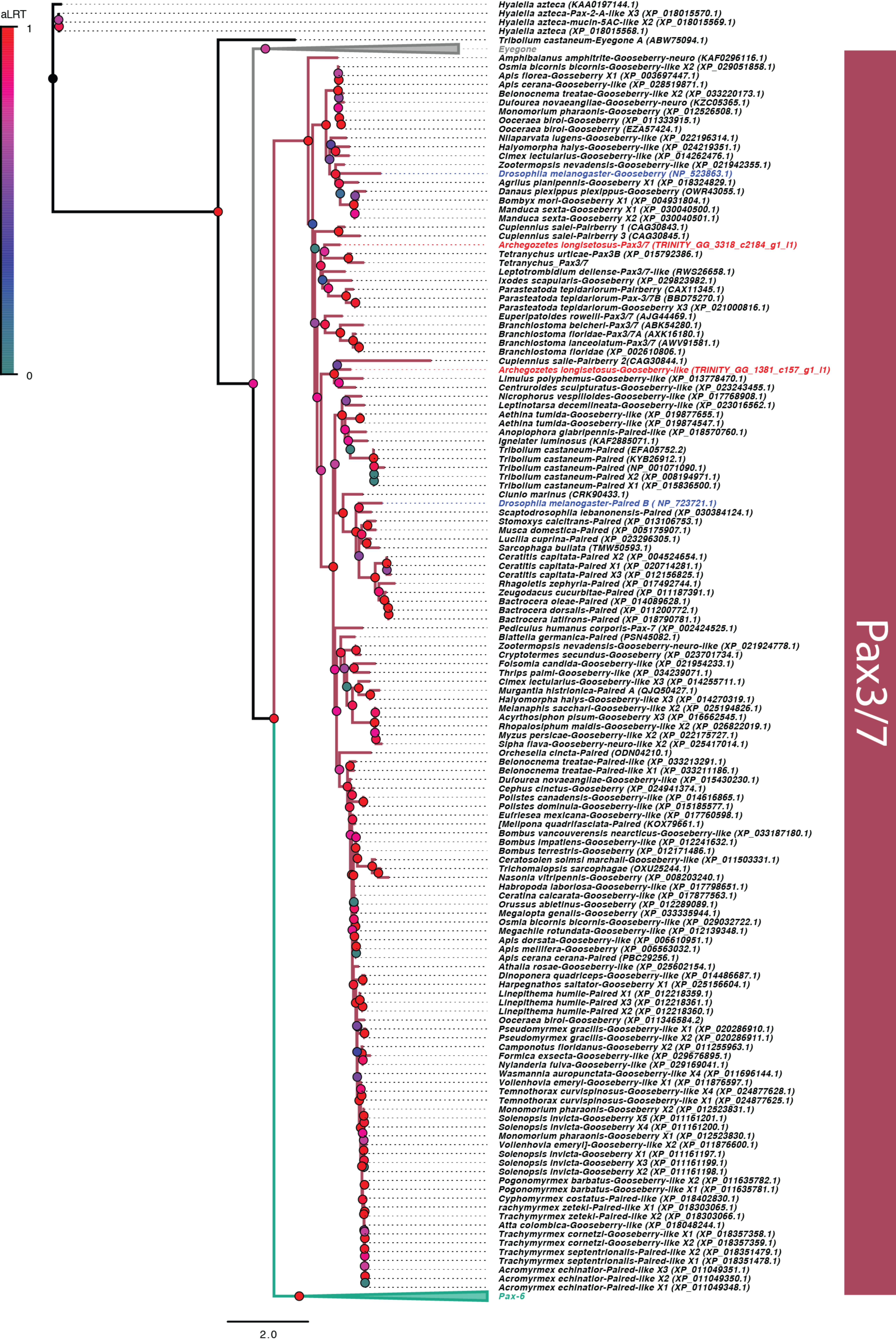

### Supplemental Figure S4D

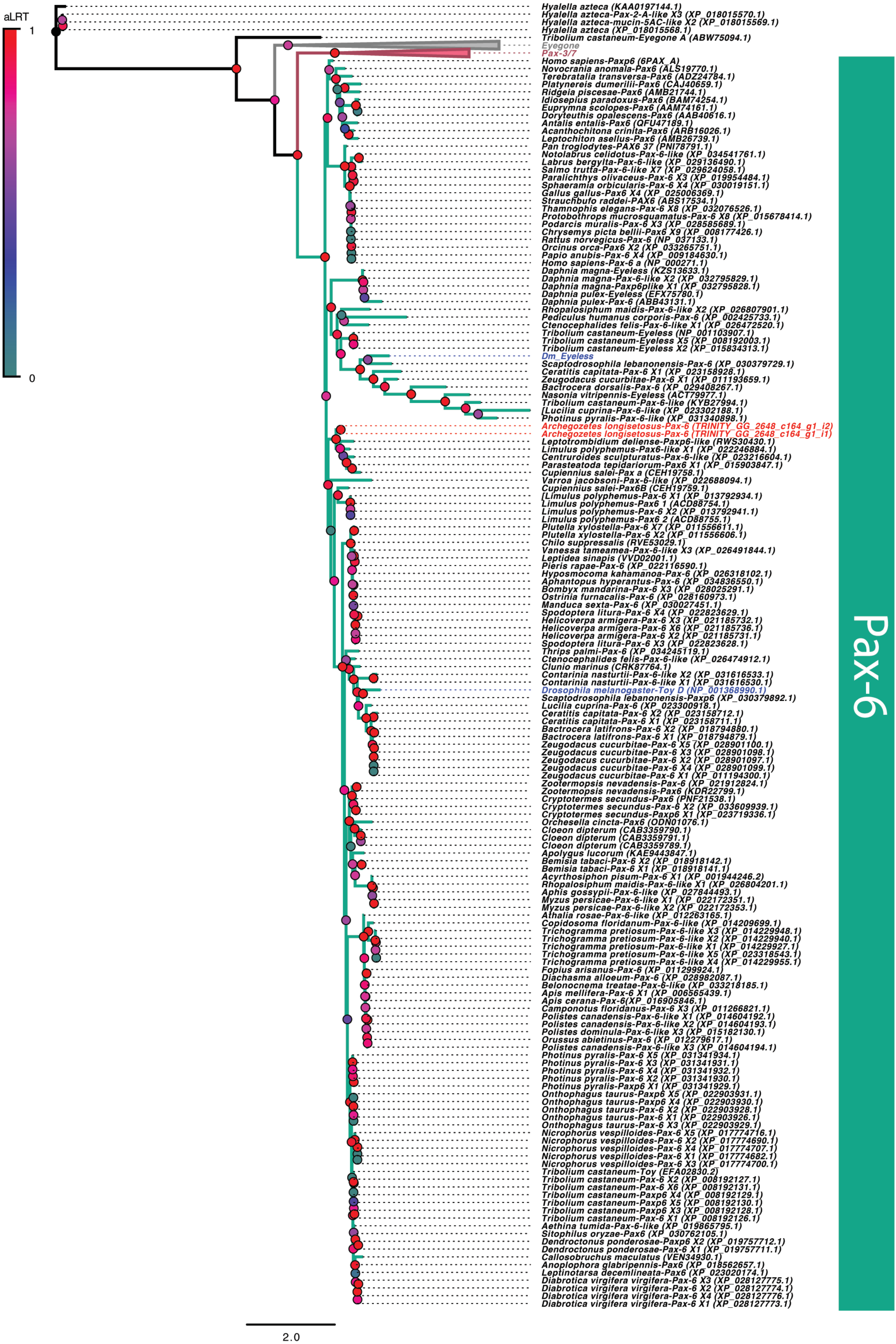

### Supplemental Figure S5

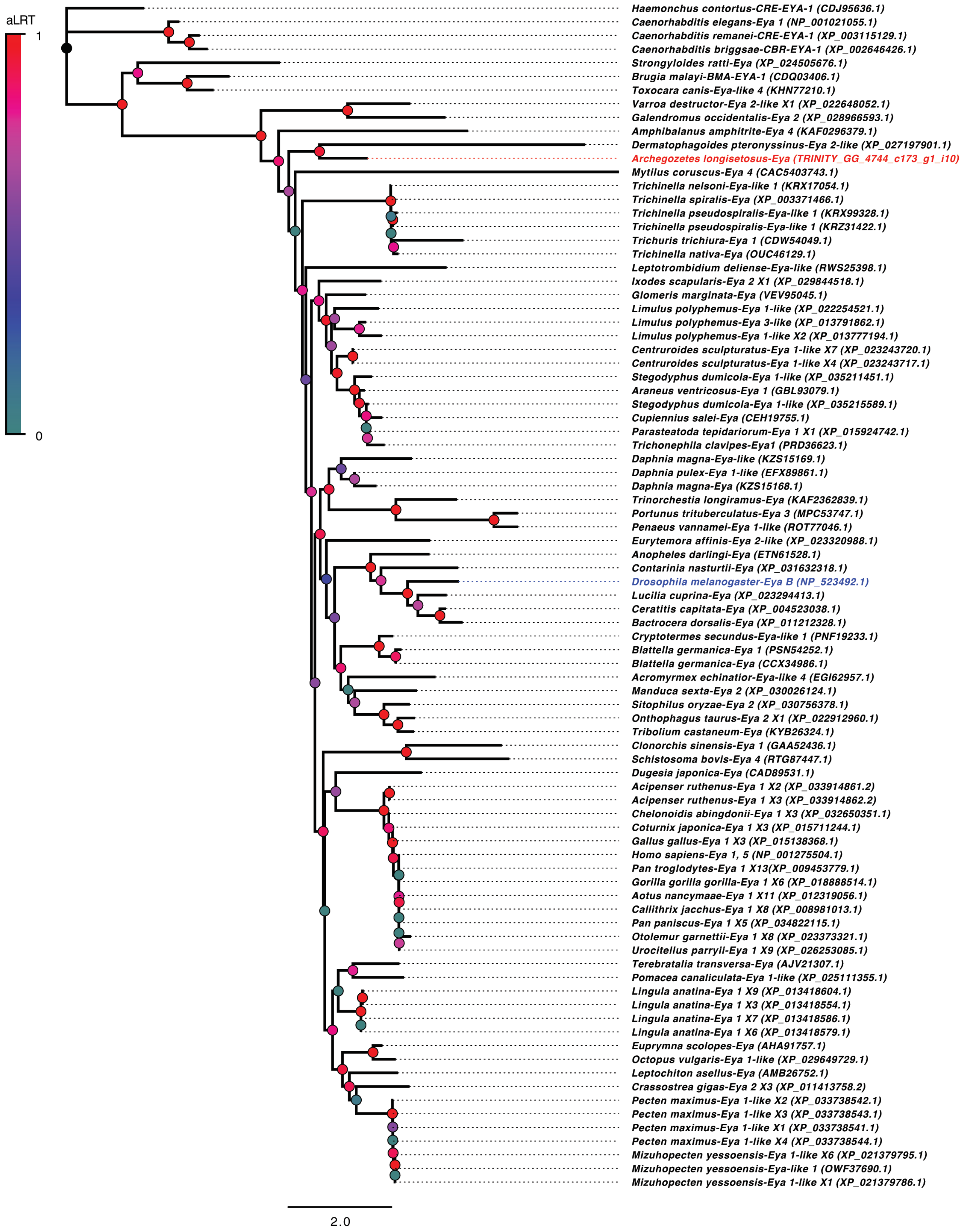

### Supplemental Figure S6

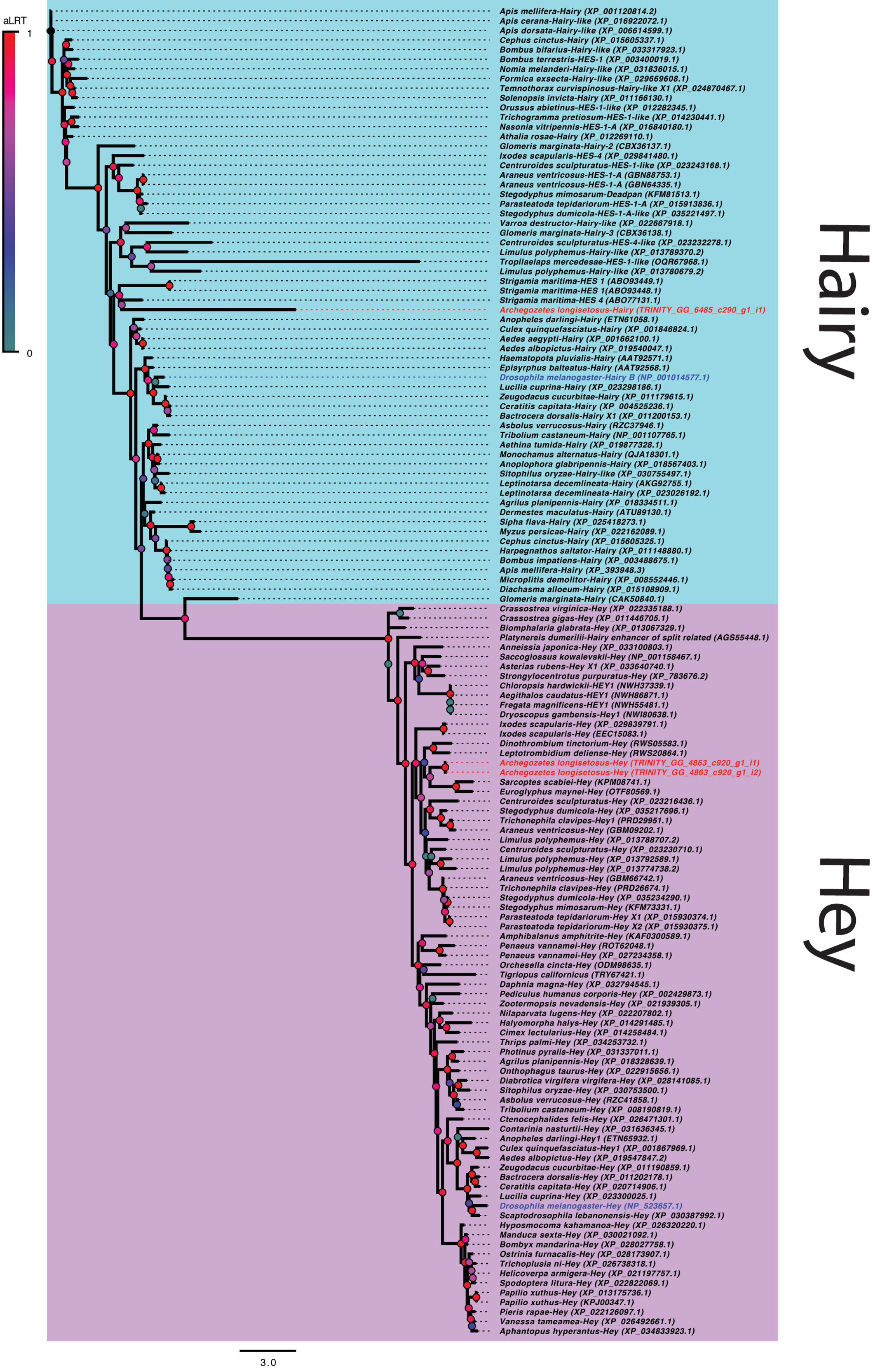

### Supplemental Figure S7

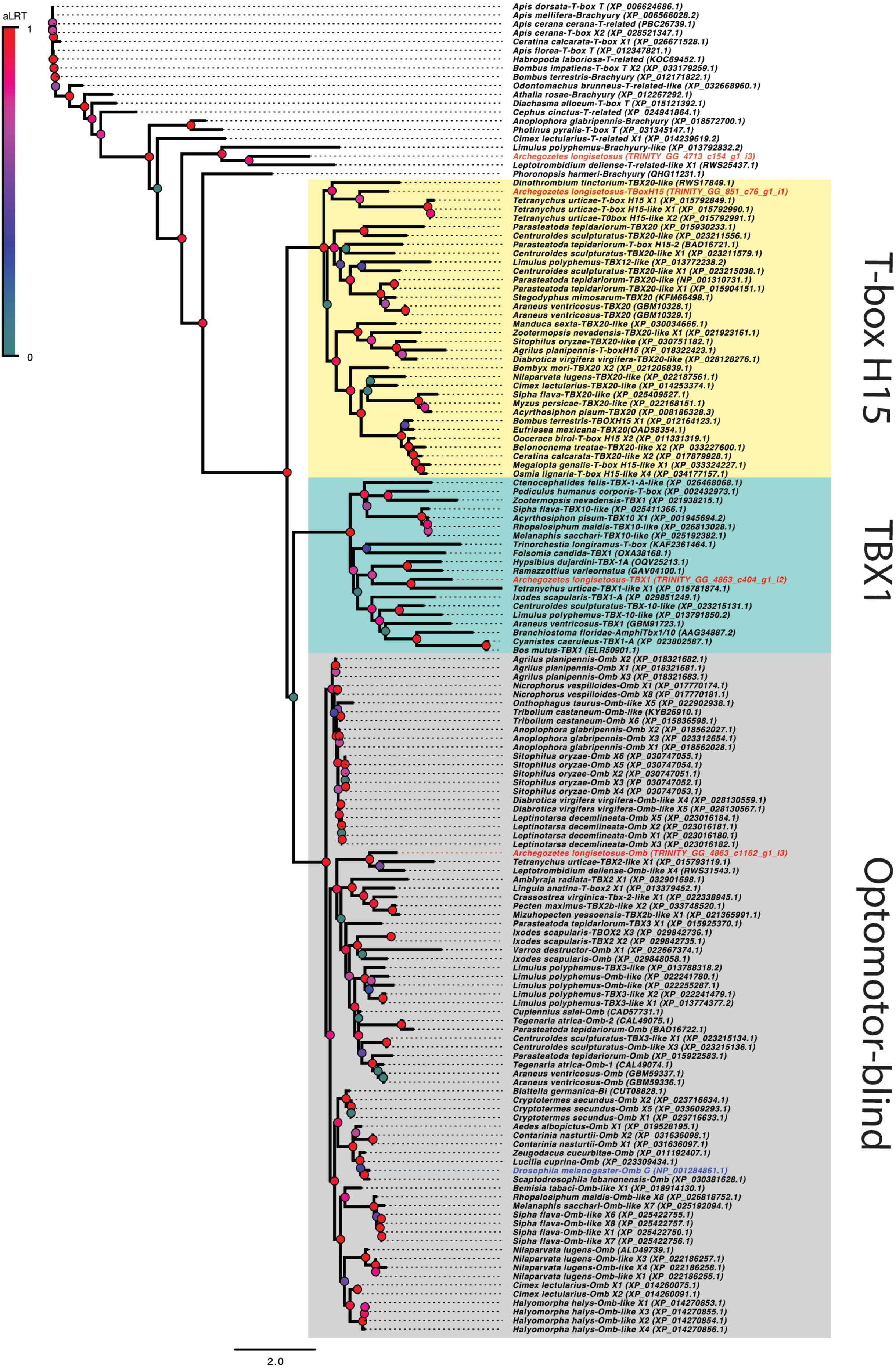

### Supplemental Figure S8

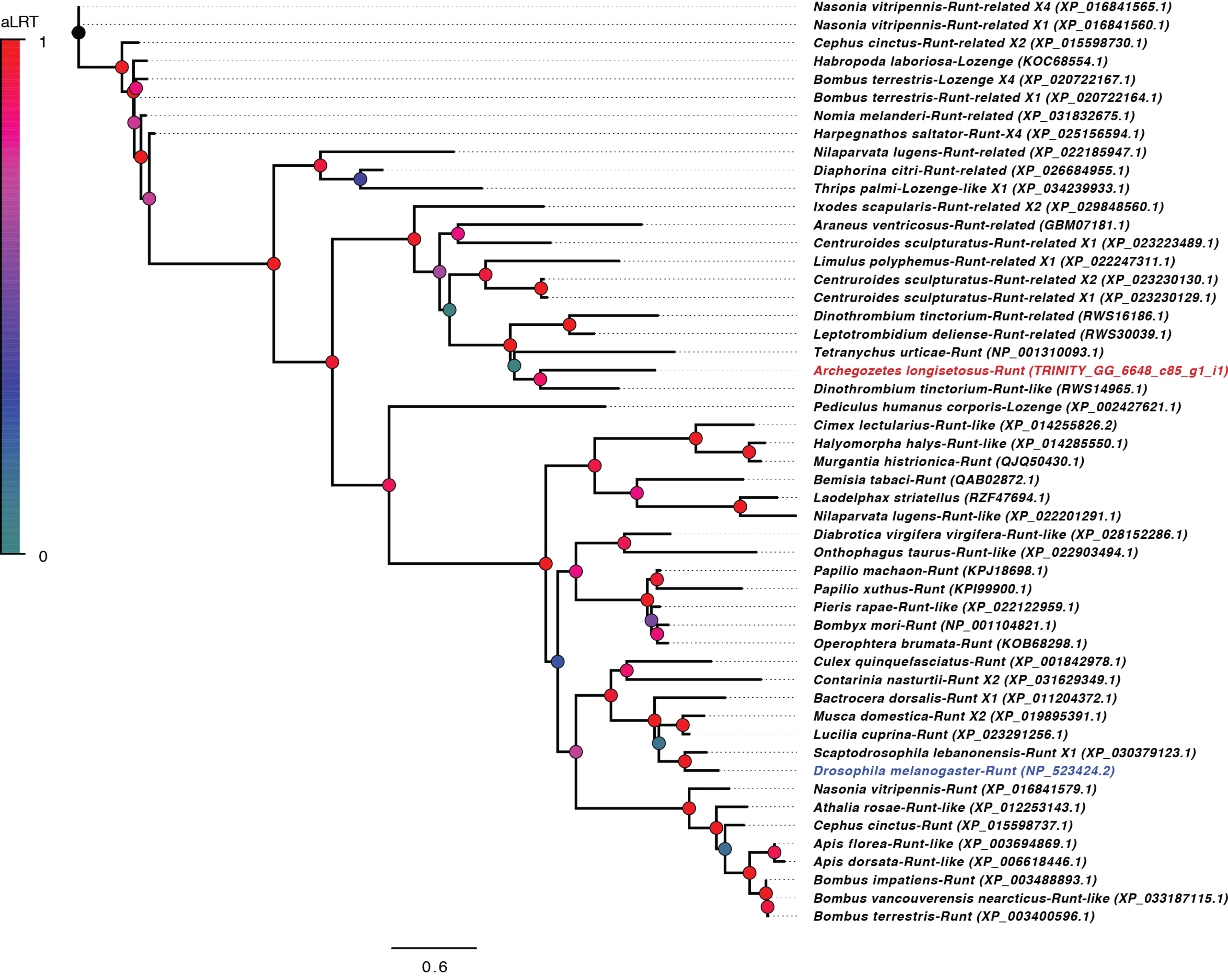

### Supplemental Figure S9

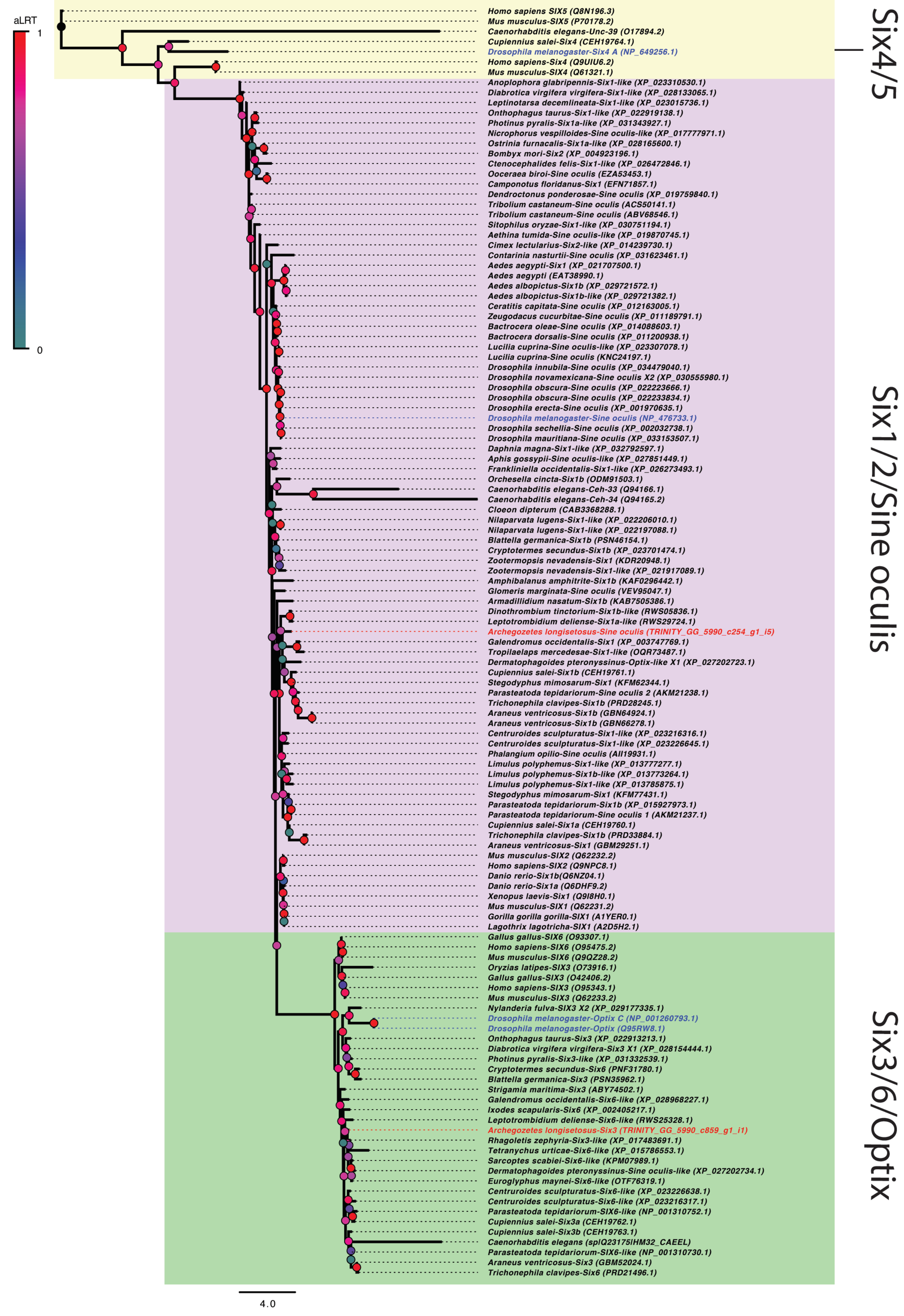

### Supplemental Figure S10

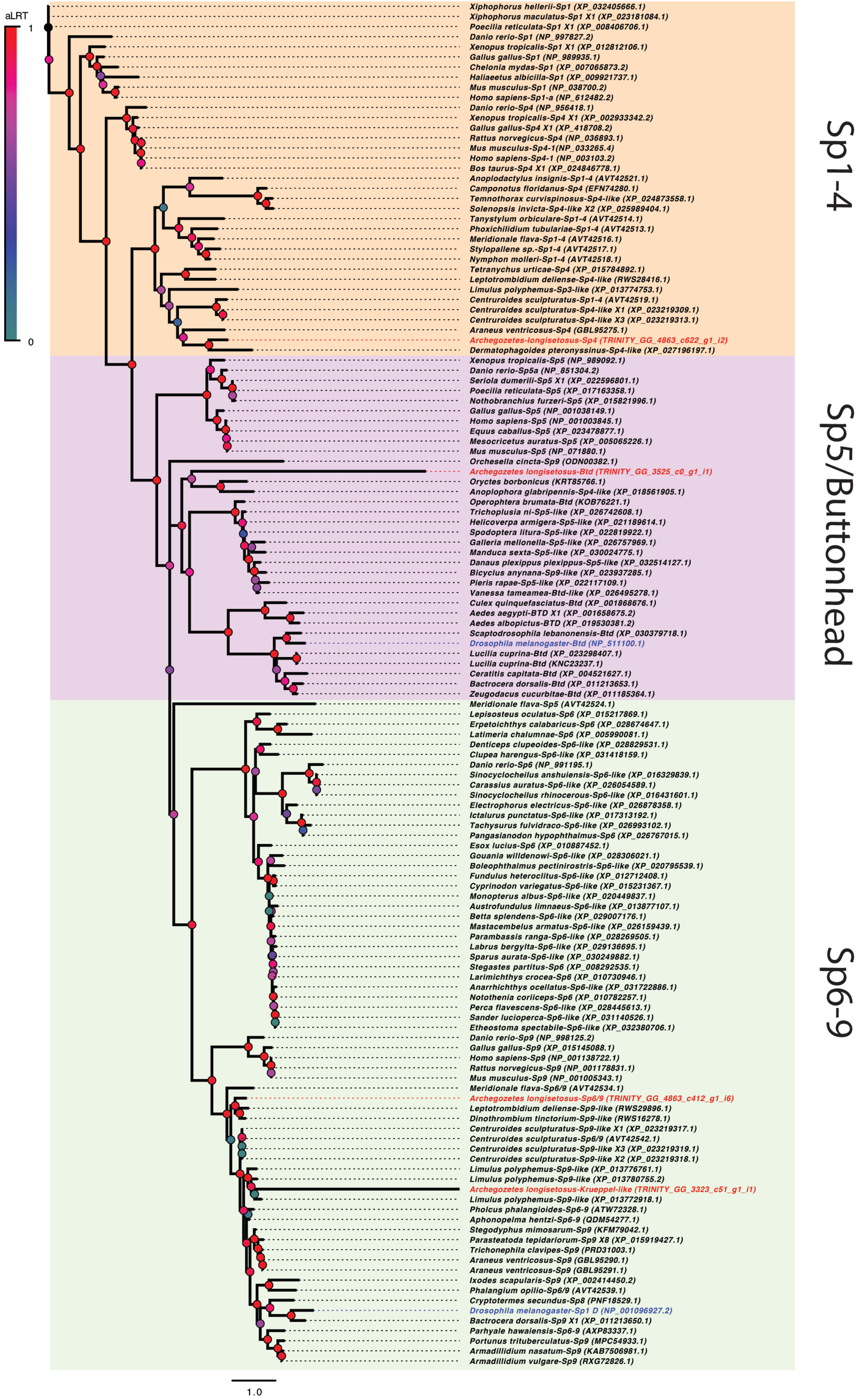

### Supplemental Figure S12

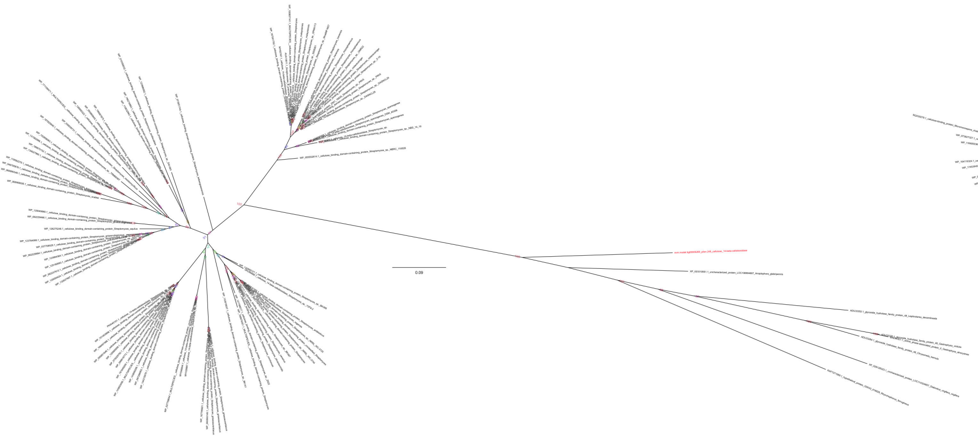

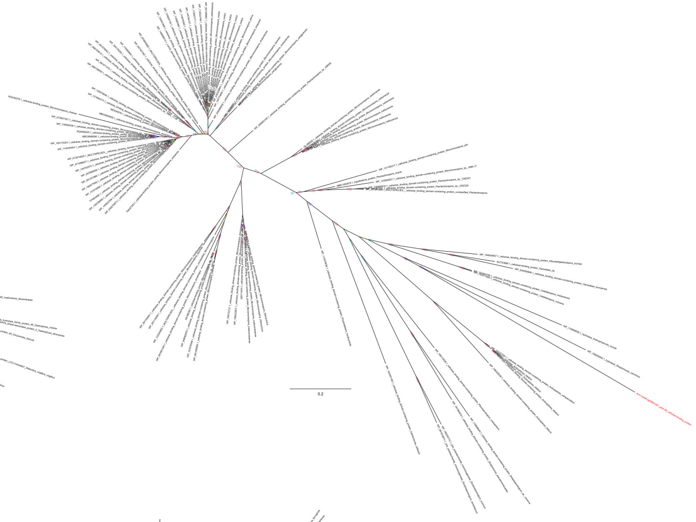

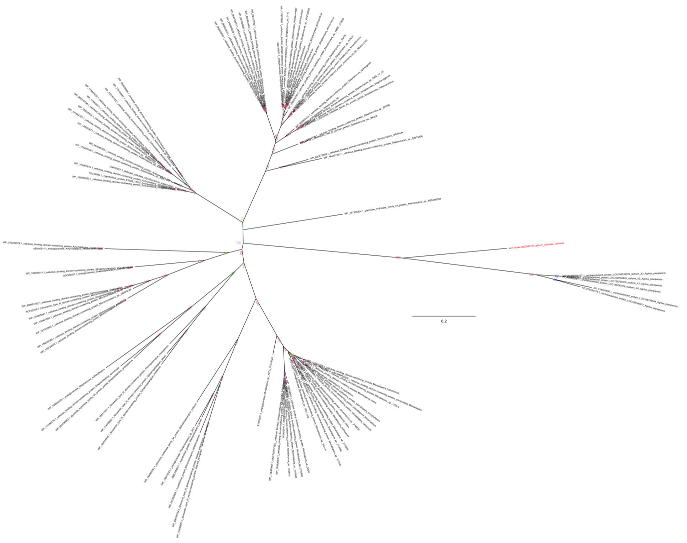

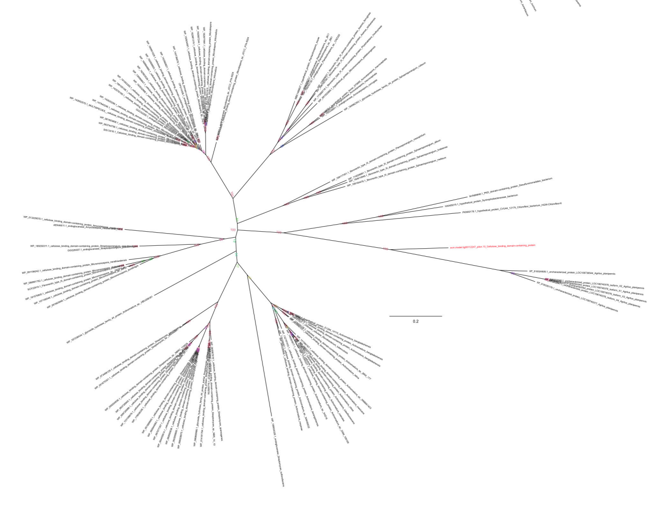

### Supplemental Figure S13

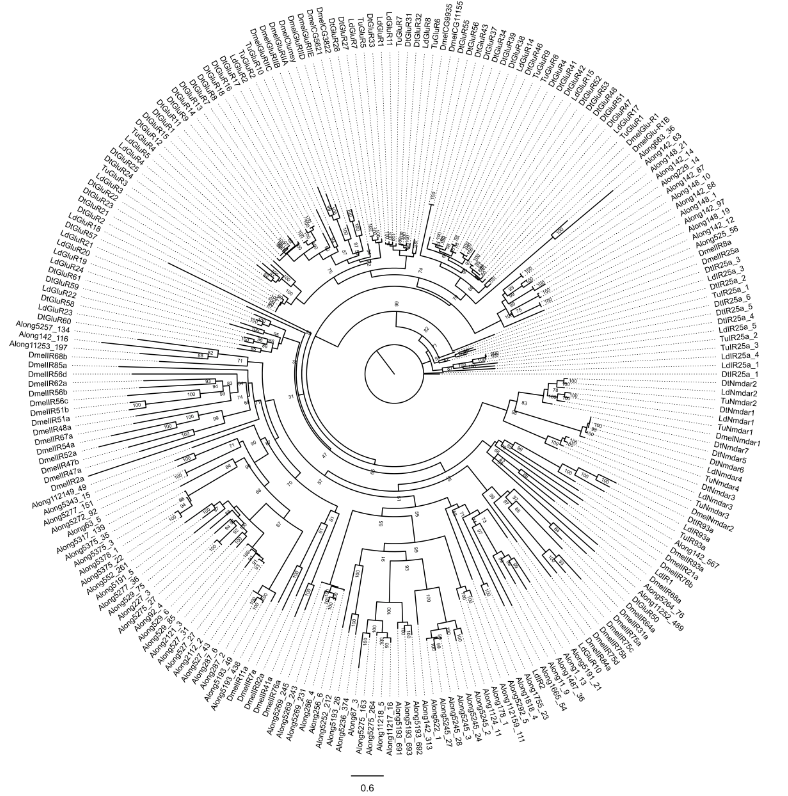

### Supplemental Figure S14

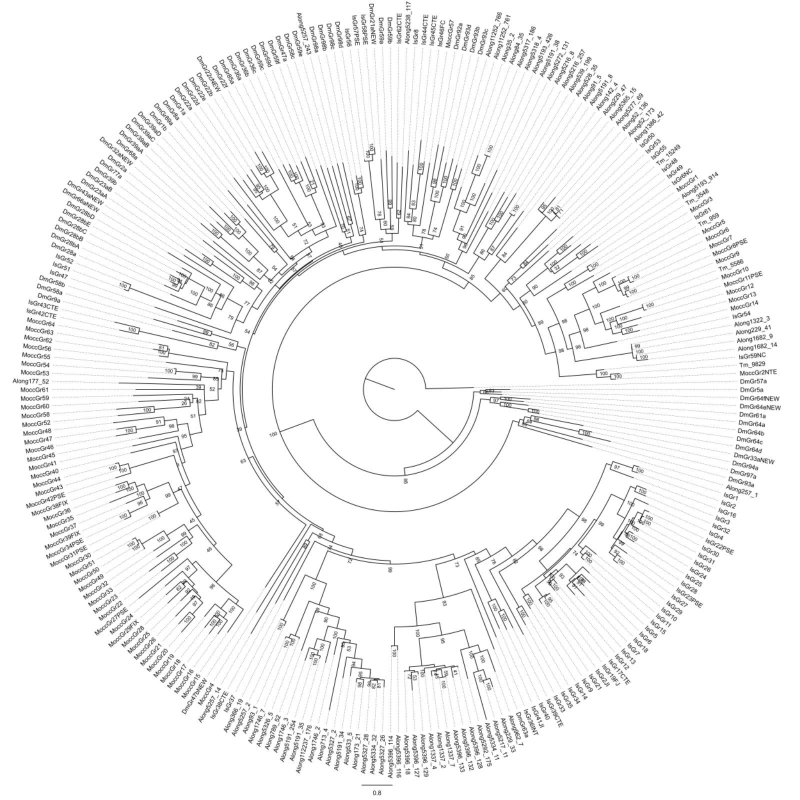
